# Stability of influenza A virus in droplets and aerosols is heightened by the presence of commensal respiratory bacteria

**DOI:** 10.1101/2024.02.05.578881

**Authors:** Shannon C. David, Aline Schaub, Céline Terrettaz, Ghislain Motos, Laura J. Costa, Daniel S. Nolan, Marta Augugliaro, Irina Glas, Marie O. Pohl, Liviana K. Klein, Beiping Luo, Nir Bluvshtein, Kalliopi Violaki, Walter Hugentobler, Ulrich K. Krieger, Thomas Peter, Silke Stertz, Athanasios Nenes, Tamar Kohn

## Abstract

Aerosol transmission remains a major challenge for the control of respiratory viruses, particularly for those that cause recurrent epidemics, like influenza A virus (IAV). These viruses are rarely expelled alone, but instead are embedded in a consortium of microorganisms that populate the respiratory tract. The impact of microbial communities and inter-pathogen interactions upon the stability of transmitted viruses is well-characterised for pathogens of the gut, but is particularly under-studied in the respiratory niche. Here, we assessed whether the presence of 5 different species of common commensal respiratory bacteria could influence the stability of IAV within droplets deposited on surfaces and within airborne aerosol particles at typical indoor air humidity. It was found that bacterial presence within stationary droplets, either a mixed community or individual strains, resulted in 10- to 100-fold more infectious IAV remaining after 1 hour. Bacterial viability was not required for this viral stabilisation, though maintained bacterial morphology seemed to be essential. Additionally, non-respiratory bacteria tested here had little stabilising effect, indicating this phenomenon was respiratory-specific. The protective bacteria stabilised IAV in droplets via induction of early efflorescence due to flattened droplet morphology during drying. Even when no efflorescence occurred at high humidity or the bacteria-induced changes in droplet morphology were abolished by aerosolization instead of deposition on a well-plate, the bacteria remained protective. This indicates an additional stabilisation mechanism that is currently unknown. Notably, respiratory bacteria at equivalent density offered varying degrees of protection in droplets, with the Gram-positive species *Staphylococcus aureus* and *Streptococcus pneumoniae* being the most robustly stabilising. This suggests that the composition of an individual’s respiratory microbiota could be a previously unconsidered host-specific factor influencing the efficacy of expelled viral spread. Identifying novel host-specific factors such as the commensal microbiota that can influence viral stability in the environment will further increase our understanding of individual transmission risks, and will provide novel opportunities to limit the spread of respiratory infections within our populations.

**Synopsis:** Our findings have significant environmental and health relevance, as they identify the host respiratory microbiota as a novel factor potentially contributing to environmental viral stability within indoor environments.

## Introduction

Influenza A virus (IAV) is a prominent respiratory pathogen that has circulated throughout the human population for over a century. Despite extensive research efforts, this virus continues to cause annual epidemics, and has caused multiple pandemics of varying scale. IAV prevalence is also seasonal, and so places recurrent pressure on healthcare systems. A major transmission pathway for this virus is via expelled particles from the cough, sneeze, or breath of an infected individual (1). These expelled plumes usually include droplets that fall and deposit on a surface within seconds (typically particles >100 µm (2)), and smaller aerosol particles that can stay airborne for minutes to hours (2). This airborne route of transmission can be extensive and is difficult to control indoors where people spend close to 90% of their time in modern society (3). Disinfection of virus-containing aerosol particles and droplets in our indoor environments is a crucial mitigation strategy to curb the spread of respiratory infections, however we have limited understanding of the processes facilitating viral survival within these particles.

Notably, viruses are not alone within the respiratory tract – the respiratory microbiota is complex and ever present, with the nasopharynx, oropharynx, and lung persistently colonised by various species of commensal bacteria. This flora is part of normal respiratory health, and bacterial colonisers typically persist with no adverse symptoms. The upper and lower respiratory tracts differ in their microbial composition (4), with common genera *Moraxella*, *Staphylococcus*, *Streptococcus, Haemophilus*, *Prevotella, Neisseria, Fusobacterium,* and *Corynebacterium* typically colonising the upper region (5–7). In contrast, the lower respiratory tract (encompassing trachea and lungs), has substantially lower biomass due to ongoing immune-mediated removal, but still harbours a number of microbes (4), with detection of *Streptococcus, Prevotella,* and *Veillonella* species reported in healthy lungs (8–10).

A particularly under-researched aspect of respiratory viral transmission is how co-expelled microbes may affect IAV stability. Airborne respiratory bacteria have been detected in expelled droplets and aerosols of saliva and sputum (11,12), and one study recently identified both *Streptococcus pneumoniae* and IAV within the same expelled aerosol fraction from co-infected ferrets Presence of bacteria was also shown to be stabilising for IAV against physical desiccation of bulk solutions, and depletion of nasal bacteria from ferrets was shown to completely abrogate aerosol-based transmission of co-infected influenza A virus from these animals (13). Presence of respiratory bacteria due to co-infection or commensal colonisation may therefore be a previously underappreciated factor of influenza virus transmission dynamics. Microbe ‘co-operation’ from the perspective of environmental stability has been investigated previously for pathogens of the gut, with binding of enteric viruses to commensal gut bacteria enhancing both viral stability and viral transmission fitness (14–16). Investigating whether similar stabilisation could occur between respiratory pathogens will improve our understanding of what factors keep these viruses infectious whilst airborne, and we seek to understand the potential mechanism(s) of pathogen stabilisation within systems representative of respiratory droplets and aerosols.

## Results

### Commensal respiratory bacteria enhance viral stability in droplets

To assess whether commensal bacteria could influence the stability of IAV in static droplets, decay experiments were performed in a humidity-controlled chamber. Here, 1-μL virus-containing droplets were deposited on a non-binding surface in relative humidity (RH) and temperature conditions that are representative of indoor spaces. In temperate climates, indoor RH can be <30% in winter months, and around 50% in summer (17), thus conditions of 40% RH at ambient temperature (22 – 25°C) were chosen as ‘mid-point’ indoor conditions. **Figure 1A** shows the decay of IAV infectivity in 1-μL droplets when the virus was deposited alone in phosphate-buffered saline (PBS). Whilst there was stability in viral titer for the first 15 minutes post-deposition, a substantial 3-log10 decay in titer was then observed after 30 minutes. After this sharp decay, remaining infectious viral titers were stabilised, giving an inverted sigmoidal decay curve over the entire 60-minute time-course. This sigmoidal decay for IAV in droplets has been observed before (18,19), with the sharpest decline in infectious titers occurring just prior to droplet efflorescence, when the salt molality of the droplets is maximal.

**Figure 1.**
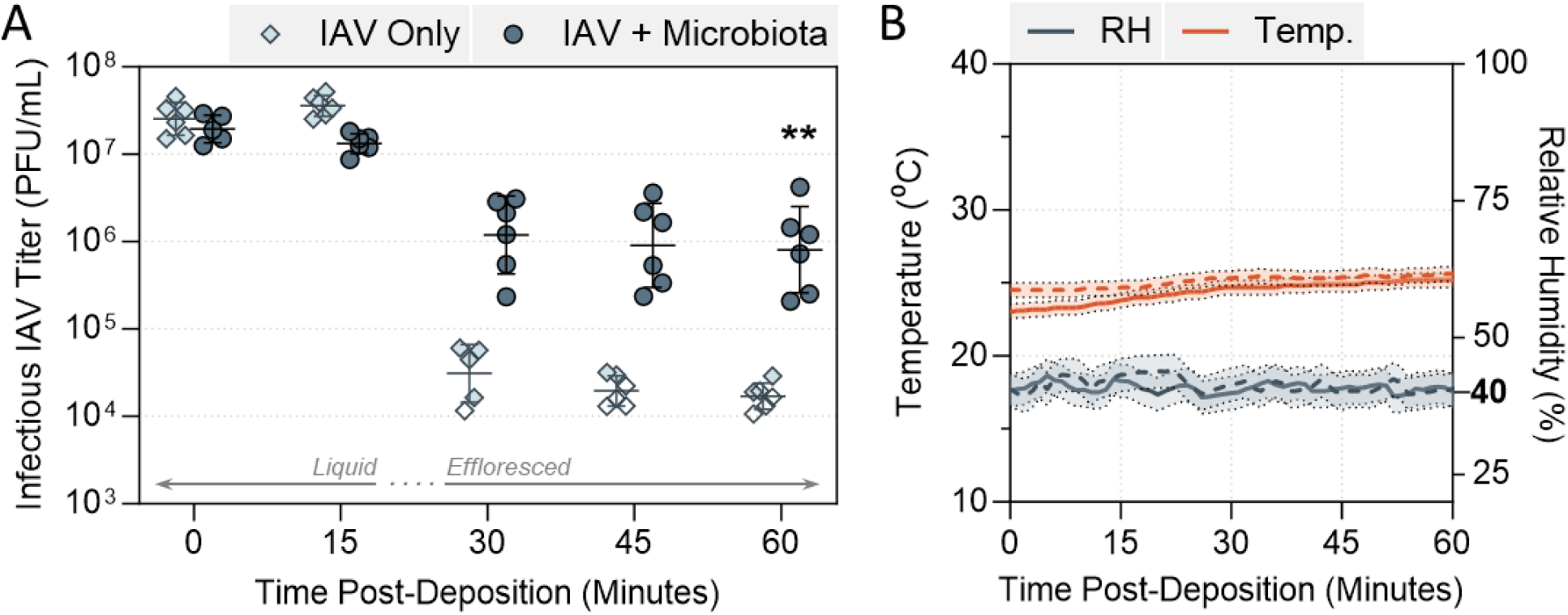
Decay of IAV in 1-μL PBS droplets at 40% relative humidity (RH) and 22 – 25°C in a humidity-controlled chamber when mixed with microbiota bacteria. **A)** Virus was added to PBS alone (IAV Only), or added to PBS containing live commensal respiratory bacteria (IAV + Microbiota), for 5 × 10^7^ PFU/mL final virus concentration. Equal CFU of *S. pneumoniae, S. aureus, H. influenzae, M. catarrhalis*, and *P. aeruginosa* were added for 10^8^ CFU/mL total in the +Microbiota case. 1-μL droplets of each virus suspension were deposited on a non-binding 96-well plate and exposed to indoor air conditions for a total of 60 minutes. Triplicate droplets of each mixture were recovered at time-points 0, 15, 30, 45, and 60 minutes post-deposition, and infectious viral titers were quantified by plaque assay (clear symbols indicate samples that were below plaque assay limit of quantification (LOQ), and were set a LOQ/√2). Infectious viral titers were corrected for physical recovery (determined by genome quantification of IAV in each recovered droplet by dPCR) relative to samples collected immediately after deposition (time 0, where no physical loss has occurred due to drying). Individual data points of triplicate droplets, from 2 independent experimental repeats are shown (n = 6 droplets total per group), with geometric mean ± geometric SD. Significant differences between IAV titers at time 60 was determined by Mann-Whitney test (**, P ≤ 0.01). **B)** RH and temperature were monitored by a portable hygrometer across the 60- minute period, with readings taken every minute. Solid and dotted lines distinguish readings from individual experimental repeats, with confidence intervals (±0.5 °C, ±3 %RH as provided by manufacturer) indicated by shaded regions. Bold text indicates the target RH.

This is in contrast to the inactivation kinetic observed when respiratory bacteria were also present within the deposited droplets. A consortium of 5 different bacteria (*Streptococcus pneumoniae, Staphylococcus aureus, Haemophilus influenzae, Moraxella catarrhalis,* and *Pseudomonas aeruginosa*, representative of the commensal microbiota in the upper respiratory tract of healthy individuals), were mixed in equal proportions and added to IAV prior to droplet deposition. This set-up was designed to mimic pathogens that may be spatially co-localised on a mucosal surface before being expelled by a cough or sneeze. This bacterial mixture was highly stabilising to the virus over the same 60-minute time-course. Approximately 1-log10 reduction in infectious viral titer was observed in this group (termed +microbiota) at 30 minutes, compared with the decay of 3-log10 for virus alone. By the end of the time-course, approximately 100-fold more infectious virus was recovered from droplets containing the microbiota mixture. In all cases, viral infectivity data was corrected for physical recovery of IAV from each individual droplet (typical work-flow of droplet experiments is shown in **Supplementary Figure S1**), hence this increased viral titer was not simply due to altered drying and reduced viral adsorption to the plate surface when bacteria were present. RH and temperature were constrained during all experiments, and were reproducible across experimental repeats (**Fig. 1B**).

### Individual Gram-positive respiratory bacteria provide viral stabilisation in droplets

To determine which of the microbiota members were most crucial for mediating protection, bacterial species were next tested individually. **Figure 2A** shows both gram-positive bacteria *S. pneumoniae* and *S. aureus* offered substantial protection to IAV against droplet-associated decay at 40% RH and 22 – 25°C (T and RH data shown in **Supp. Fig. S2A**). Over the entire monitoring period of 60 minutes, only 1-log10 decrease in viral titer was observed when mixed with each of these individual bacteria, compared to >3-log10 decay for IAV alone. This was very similar to the trend seen in **Figure 1A**, showing individual bacteria can be just as effective as the mixed microbiota for viral stabilisation. Again, virus alone decayed most rapidly between the 15 – 30-minute time-points, and this sharp titer decrease was mitigated by the presence of bacteria. Additionally, the drying behaviour of the droplets was vastly different when bacteria were present; droplets containing bacteria effloresced sooner (first visible instance of bacteria-containing droplet efflorescence is indicated by light and dark green circles in **Fig. 2B**), and showed a wider/flatter morphology compared to droplets containing virus alone. Diameters of the drying droplets were quantified using an automated particle identification software (with a custom model pre-trained on droplet images) paired with the particle measurement function in ImageJ, and diameter measurements of drying droplets are shown in **Supplementary Figure S3A**.

**Figure 2.**
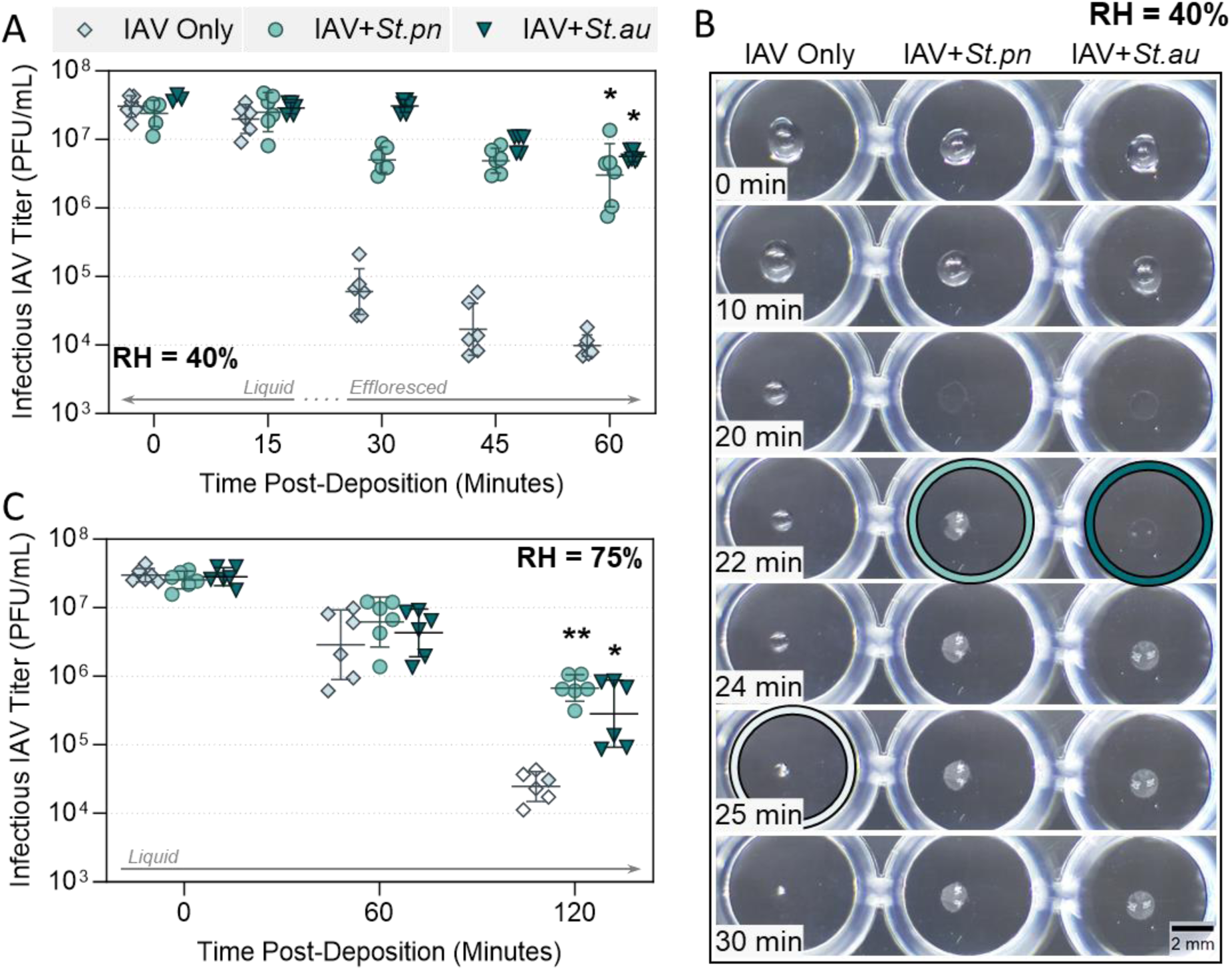
Decay of IAV in 1 µL PBS droplets at 22 – 25°C in a humidity controlled chamber when mixed with Gram-Positive bacteria. **A)** Virus was added to PBS alone (IAV Only), or added to PBS containing live *Streptococcus pneumoniae* (+*St.pn*) or *Staphylococcus aureus* (+*St.au*) bacteria at 10^8^ CFU/mL. In all cases, virus was added for 5 × 10^7^ PFU/mL final concentration. 1µL droplets of each virus suspension were deposited on a non-binding 96-well plate and exposed to indoor air conditions for a total of 60 minutes at 40% RH. Triplicate droplets of each mixture were recovered at time-points 0, 15, 30, 45, and 60 minutes post-deposition, and infectious viral titers were quantified by plaque assay. Infectious viral titers were corrected for physical recovery (determined by genome quantification of IAV in each recovered droplet by dPCR) relative to samples collected immediately after deposition (time 0, where no physical loss has occurred due to drying). Individual data points of triplicate droplets from 2 independent experimental repeats are presented (n = 6 droplets total per group), with geometric mean ± geometric SD. Significant differences in infectious titers at time 60 relative to IAV only control were determined by One-Way ANOVA (*, P ≤ 0.05). **B)** 1-µL droplets containing IAV alone or IAV mixed with *S. pneumoniae* or *S. aureus* in PBS [as in (A)] were recorded during drying at 40% RH. Images are representative of 3 individual droplets per group. Coloured circles highlight the time-point where crystallisation was first visible by eye for each sample. Scale bar = 2 millimetres (mm) **C)** Decay of IAV in 1-µL droplets at 75% RH, when IAV was added to PBS alone, or to PBS containing *S. pneumoniae* or *S. aureus* [as in (A)]. 1-µL droplets were deposited on a non-binding 96-well plate and exposed to indoor air at 75% RH for a total of 120 minutes. Triplicate droplets of each mixture were recovered at time-points 0, 60, and 120 minutes post-deposition. Infectious titers quantified and corrected as for (A), clear symbols indicate samples that were below plaque assay limit of quantification (LOQ), and were set a LOQ/√2 as described in Methods. Individual data points of triplicate droplets from 2 independent experimental repeats (n = 6 droplets total per group) are presented, with geometric mean ± geometric SD. Significant differences in infectious titers at time 120 relative to IAV only control were determined by Kruskal-Wallis test (*, P ≤ 0.05, **, P ≤ 0.01).

To ensure that the observations conducted with lab adapted A/WSN/33 IAV were also applicable to clinically relevant strains, a comparison was made to A/Netherlands/07/2009, a more recent human isolate of H1N1 IAV. **Supplementary Figure S4** shows the inactivation kinetic of the virus alone in droplets was comparable between these two IAV strains, and that similar stabilisation by *S. pneumoniae* occurred for both. A/WSN/33 was therefore deemed a suitable surrogate, and was used for the entirety of this study given the ease of high titer growing and viral titration. Literature has reported that certain bacteria can improve viral binding to target host cells by acting as ‘coupling agents’, and thus before progressing further it was important to ensure enhanced viral titers in droplet experiments were not simply due to bacterial effects in the down-stream plaque assays. **Supplementary Figure S5** showed that cells exposed to virus and bacteria together showed the same levels of virus infection compared to virus alone at ratios representative of those used in droplet experiments, indicating that titre differences in these experiments were due to a true stabilising effect of the bacteria.

We know from prior droplet work (19) that IAV inactivation kinetics differ between efflorescing and deliquesced conditions, hence it was of interest to assess whether bacteria could also be protective within a deliquesced droplet. Droplet experiments were performed at a higher RH of 75% and 22 – 25°C (T and RH data shown in **Supp. Fig. S2B**), which retained the droplets in a liquid state for the entire monitoring period (the exposure time was also increased to 2 hours, as droplet evaporation was now much slower). **Figure 2C** shows droplets containing IAV only displayed ∼3-log10 decrease in viral infectivity under these altered conditions. Over the first hour, a small amount of viral decay was seen across all samples (∼1-log10), and subsequently, a >2 log10 loss in IAV titer occurred between time-points 60 and 120 for virus alone. *S. pneumoniae* and *S. aureus* both retained the ability to protect IAV from decay at this high RH, with approximately 1-log10 more infectious virus detected in these groups at the end of the time-course.

### Individual Gram-negative respiratory bacteria provide a milder stabilisation effect for IAV in droplets compared to Gram-positive strains

Droplet experiments were repeated at 40% and 75% RH, now using the Gram-negative respiratory bacteria of interest (*H. influenzae, M. catarrhalis,* and *P. aeruginosa*) to assess whether these individual species could also be stabilising. **Figure 3A** shows the inactivation kinetics of virus alone compared to virus co-deposited with these 3 individual bacteria at 40% RH. The inverted sigmoidal inactivation curve for virus alone was again apparent, and in the presence of Gram-negative bacteria, the virus decay curves now also showed a drop in viral titer around the efflorescence point. However, Gram-negative bacteria still retained approximately 1 – 1.5-log10 higher viral titers at the end of the time-course compared to virus alone (note that IAV titres in the +*P. aeruginosa* experiment were not statistically higher than the IAV only control). Representative images in **Figure 3B** show that these Gram-negative bacteria had a similar effect on the physical behaviour of the droplets compared with Gram-positive bacteria, causing an earlier efflorescence time, paired with a wider/flatter morphology and larger final radius (see **Supplementary Figure S3B** for diameter measurements). When tested at the higher humidity of 75% RH, all three Gram-negative strains retained the ability to stabilise IAV (**Fig. 3C**), though IAV titres in the +*H. influenzae* experiment were no longer statistically higher than controls. The most consistently stabilising Gram-negative bacterium was *M. catarrhalis,* which at 40% RH also caused earliest efflorescence and retained the widest droplet diameter.

**Figure 3.**
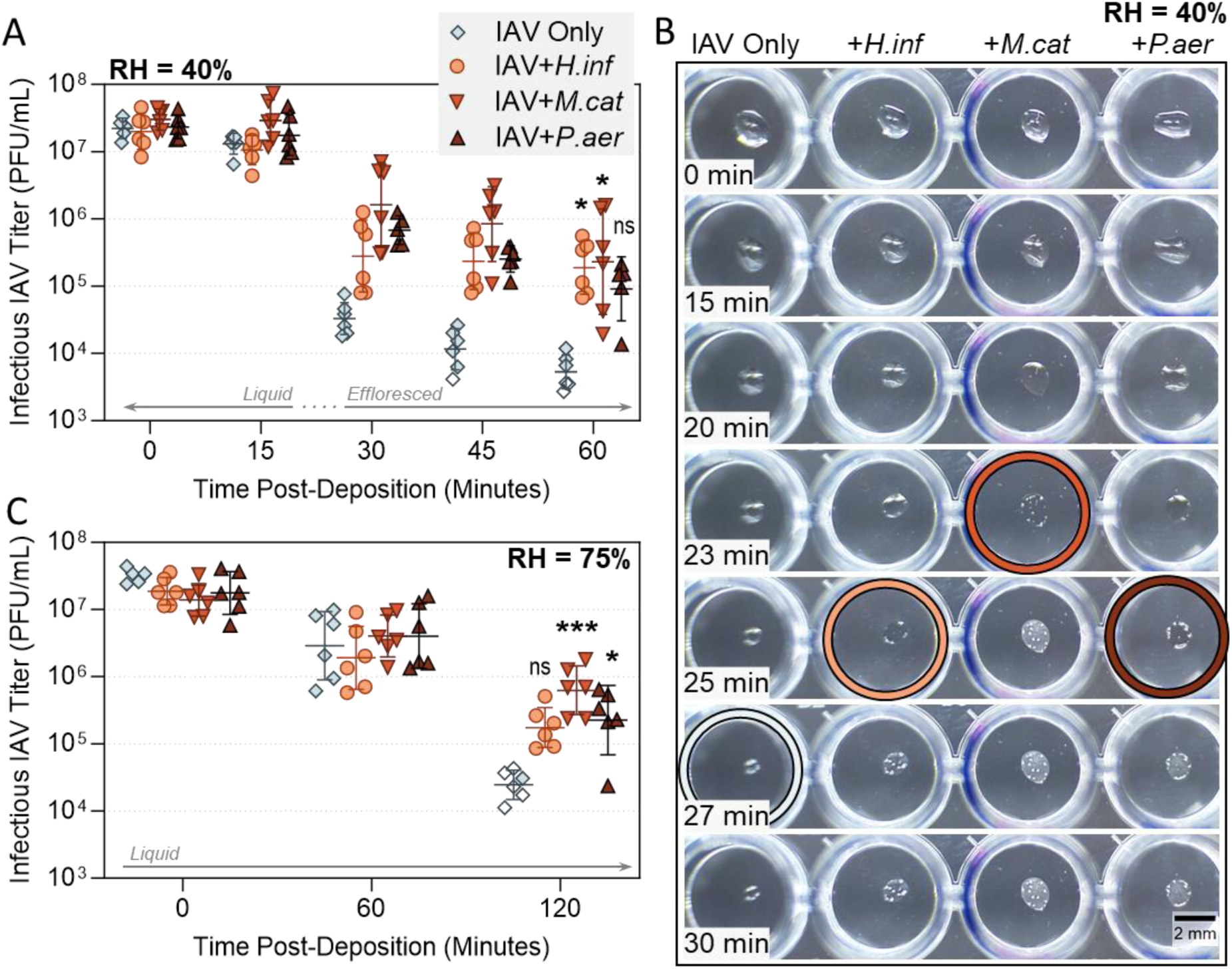
Decay of IAV in 1-µL PBS droplets at 22 – 25°C in a humidity controlled chamber when mixed with Gram-Negative bacteria. **A)** Virus was added to PBS alone (IAV Only), or added to PBS containing live *Haemophilus influenzae* (+*H.inf*), *Moraxella catarrhalis* (+*M.cat*), or *Pseudomonas aeruginosa* (+*P.aer*) bacteria at 10^8^ CFU/mL final concentration. In all cases, virus was added for 5 × 10^7^ PFU/mL final concentration. 1-µL droplets of each virus suspension were deposited on a non-binding 96-well plate and exposed to indoor air conditions for a total of 60 minutes at 40% RH. Triplicate droplets of each mixture were recovered at time-points 0, 15, 30, 45, and 60 minutes post-deposition, and infectious viral titers were quantified by plaque assay (clear symbols indicate samples that were below plaque assay limit of quantification (LOQ), and were set a LOQ/√2). Infectious viral titers were corrected for physical recovery (determined by genome quantification of IAV in each recovered droplet by dPCR) relative to samples collected immediately after deposition (time 0, where no physical loss has occurred due to drying). Data shows individual data points of triplicate droplets, from 2 independent experimental repeats (n = 6 droplets total per group), with geometric mean ± geometric SD. Significant differences in infectious titers at time 60 relative to IAV only control were determined by Kruskal-Wallis test (*, P ≤ 0.05, ns, not significant). **B)** 1-µL droplets containing IAV alone or IAV mixed with *H. influenzae, M. catarrhalis*, or *P. aeruginosa* in PBS [as in (A)] were recorded during drying at 40% RH. Images are representative of 3 individual droplets per group. Coloured circles highlight the time-point where crystallisation was first visible by eye for each sample. Scale bar = 2 millimetres (mm). **C)** Decay of IAV in 1-µL droplets at 75% RH, when IAV was added to PBS alone, or to PBS containing *H. influenzae, M. catarrhalis*, or *P. aeruginosa* [as in (A)]. 1-µL droplets were deposited on a non-binding 96-well plate and exposed to air at 75% RH for a total of 120 minutes. Triplicate droplets of each mixture were recovered at time-points 0, 60, and 120 minutes post-deposition. Infectious titers were quantified and corrected as for (A), clear symbols indicate samples that were below plaque assay limit of quantification (LOQ), and were set a LOQ/√2). Individual data points of triplicate droplets from 2 independent experimental repeats (n = 6 droplets total per group) are presented, with geometric mean ± geometric SD. Significant differences in infectious titers at time 120 relative to IAV only control were determined by Kruskal-Wallis test (*, P ≤ 0.05, ***, P ≤ 0.001, ns, not significant).

### Bacteria do not affect the efflorescence relative humidity, but rather alter the droplet morphology for faster evaporation

To test whether the observed bacterial-induced morphological differences between droplets also affected the evaporation kinetics, we performed fast-drying experiments similar to those of **Figures 2** and **3**. First, 1-μL droplets were deposited on a hydrophobic cover slip, then placed in an environmental cell with precision controlled temperature and relative humidity. These kinetic tests were conducted in a non-Biosafety Level 2 laboratory, thus no virus was included and heat-inactivated bacteria were used for biosafety purposes. **Figure 4A** shows exposure of 1-μL droplets (PBS only, or PBS+*S. pneumoniae* bacteria) to an initially high humidity (>85% RH) with subsequent fast switching to an RH of ∼40%. The RH progression between these fast-drying experiments was identical, shown by the overlapping continuous lines. Significantly delayed efflorescence (3.57 ± 0.16 minutes, P = 0.0232, paired two-tailed t-test) was also observed at 40% RH for droplets not containing any bacteria, consistent with observations in **Figures 2B** and **3B** which used live bacteria and IAV. During these controlled humidity cycles at constant temperature, morphological changes of the droplets were also monitored optically with a microscope to acquire images and movies of the droplet morphology. **Supplementary Videos S1** and **S2** show kinetically slower evaporation for 1-μL PBS only droplets with a half sphere geometry, compared to PBS+bacteria droplets which exhibited a more “pancake” like morphology during drying. As the latter has a larger surface area exposed to the dry atmosphere, this would result in a kinetically faster evaporation and time shift in efflorescence relative to PBS alone.

**Figure 4.**
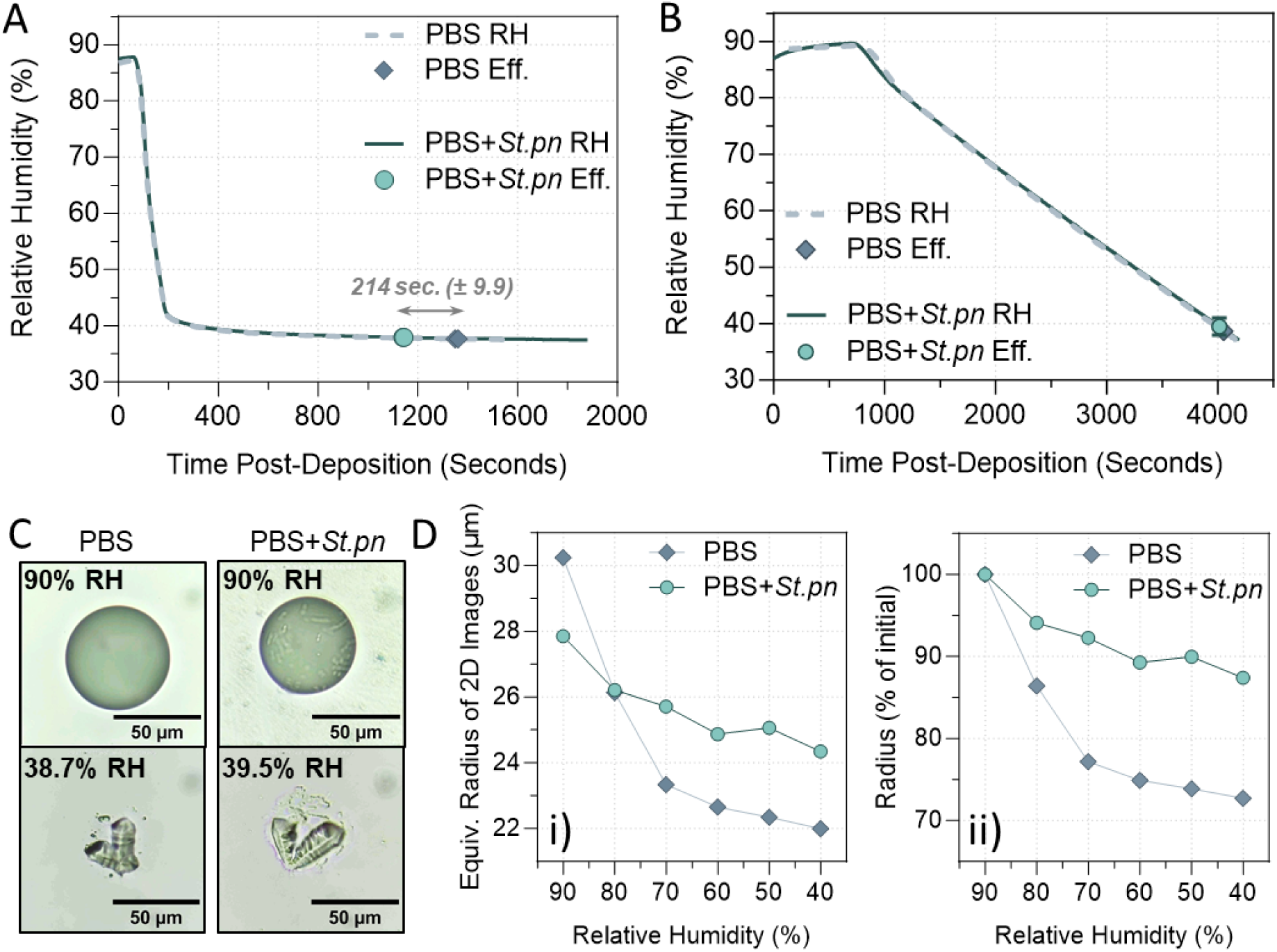
Confirmation of altered efflorescence behaviour by whole bacterial cells. **A)** RH versus time for two rapid drying experiments with large droplets (1 μL initial volume) of PBS alone or PBS containing heat-inactivated *S. pneumoniae* at 10^8^ CFU equivalent/mL (PBS+*St.pn*). Thick lines indicate RH ramp for each independent experiment (PBS: grey dashed line, PBS+*St.pn:* green solid line), whilst symbols indicate the exact time when efflorescence (Eff.) occurred for both droplet types. PBS droplets without bacteria effloresced an average of 214 ± 9.9 seconds later than the corresponding droplets containing bacteria (n = 2 experiments per group). **B)** RH versus time for slow drying experiments, using smaller printed droplets and a drying rate of ca.1.4·10^-2^ %RH/s. Drying data is shown for two droplets; thick lines indicate RH ramp for each independent experiment (PBS: grey dashed line, PBS+*St.pn:* green solid line), whilst symbols indicate the efflorescence relative humidity (ERH) for each condition. Both ERH agreed within measurement uncertainty (ERH = 38.7±1.5 % for PBS, n = 4 individual experiments; and ERH = 39.5±1.5% RH for PBS+*St.pn*, n = 3 individual experiments) **C)** Representative images of each droplet type at the indicated RH during the slow drying experiment of [B]. Scale bar = 50 μm. **D)** Equivalent circle radii of the 2D images for the droplets of [B] upon slow drying, presented as i) the measured 2D radii in μm, and ii) the 2D radii as a percentage of the initial radius measured at 90% RH.

To specifically measure efflorescence relative humidity (ERH), additional experiments were performed using ∼30-micron size, single printed droplets of PBS alone or PBS+bacteria. Briefly, these droplets were deposited with a droplet-on-demand printer onto a hydrophobic cover slip, and again were placed in the environmental cell. This time, RH was slowly ramped from high RH (>85%) to ∼40% over the course of an hour, allowing the droplets to continuously equilibrate with the surrounding RH. Here we observed no significant difference in ERH for droplets containing bacteria compared to those without bacteria (**Fig. 4B** and **4C**). These slow-ramp experiments were repeated 6 times with varying drying rates, with the same result (also see **Supplementary Videos S3** and **S4** for filmed printed droplets during slow-ramp drying). This indicates that the bacteria were not acting as internal nucleation cores to trigger a higher ERH, but rather, they induced faster evaporation solely by altering the droplet drying morphology.

More quantitatively, differences in droplet morphology become evident upon measuring the equivalent circle radius of the droplet 2D images acquired during drying. Data is shown as measured radii for each droplet with varying RH (**Fig. 4Di**) and as the percentage of the initial measured radius (**Fig. 4Dii**) during slow ramp experiments. Evidently, we observed a significantly stronger dependence of the equivalent 2D circle radius on humidity for the PBS only droplet compared with the droplet containing bacteria. While the equivalent radius of the PBS only droplet shrunk by a factor of 1.38 upon changing the relative humidity from 90% to 40%, the droplet containing bacteria shrunk by a factor of 1.14 only, indicative of a smaller contact angle of this droplet compared to the one of PBS. As hygroscopic growth for both droplets should be essentially the same, given by the water activity of PBS, these differences in equivalent radii are due to the different morphologies of both droplets, with the PBS droplet exhibiting a shape closer to a half sphere, while the droplet with bacteria flattened upon drying with much smaller change in 2D equivalent radius.

### Intact respiratory bacterial cells are required for viral stabilisation in droplets

To confirm whether the observed viral stabilisation was restricted to bacteria, or if any large inclusion could offer stabilisation to IAV in droplets, inorganic polystyrene beads of a similar size as bacteria (∼1 μm average diameter) were tested. Contrary to organic bacterial cells, these beads appeared to have no stabilising effect on IAV when co-deposited in droplets (**Fig. 5A**). Droplet filming showed that these polystyrene beads had minimal effect on the drying behaviour of the droplets compared to those containing IAV alone, with both droplet types retaining a spherical morphology during drying, and efflorescing at the same time-point (**Fig. 5B**).

**Figure 5.**
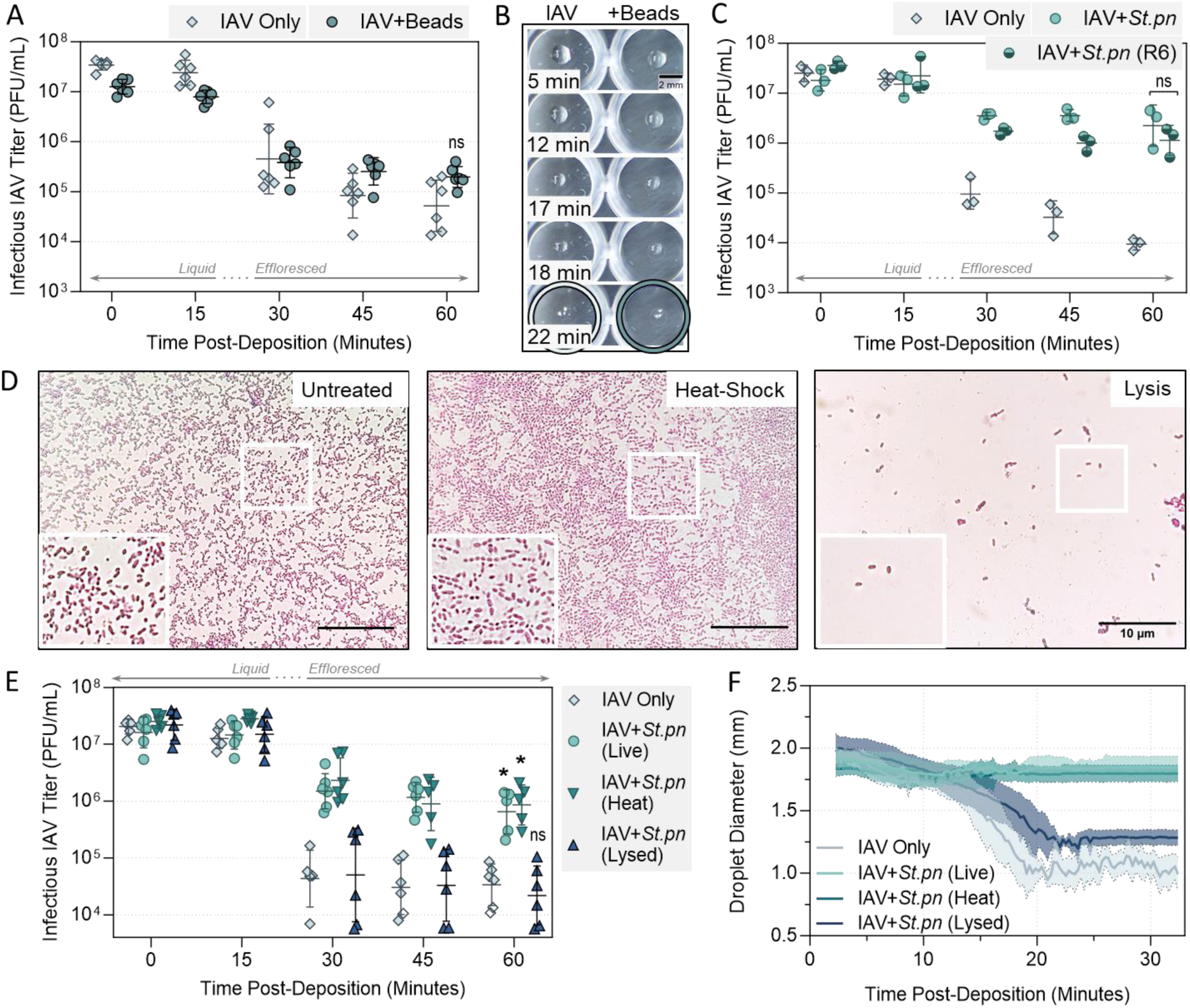
Decay of IAV in 1-µL PBS droplets at 22 – 25°C in a humidity controlled chamber when mixed with physical variations of *S. pneumoniae* bacteria. **A)** Virus was added to PBS alone (IAV Only), or added to PBS containing bacteria-like inert polystyrene 1-µm beads (IAV+beads) at 10^8^ particles/mL. In all cases, virus was added for 5 × 10^7^ PFU/mL final concentration. 1-µL droplets of each virus suspension were deposited on a non-binding 96-well plate and exposed to indoor air conditions for a total of 60 minutes at 40% RH. Triplicate droplets of each mixture were recovered at time-points 0, 15, 30, 45, and 60 minutes post-deposition, and infectious viral titers were quantified by plaque assay. Infectious viral titers were corrected for physical recovery (determined by genome quantification of IAV in each recovered droplet by dPCR) relative to samples collected immediately after deposition (time 0, where no physical loss has occurred due to drying). Data points of triplicate droplets are shown, from 2 independent experimental repeats (n = 6 droplets total per group), presented as geometric mean ± geometric SD. Significant differences in infectious titers at time 60 relative to the IAV only control were determined by unpaired t-test (ns, not significant). **B)** 1-µL droplets of PBS containing IAV alone or IAV mixed with polystyrene latex spheres (+beads) [as in (A)] were recorded during drying at 40% RH. Images are representative of 3 individual droplets per group. Coloured circles highlight the time-point where crystallisation was first visible by eye for each sample. Scale bar = 2 millimetres (mm). **C)** Decay of IAV in 1-µL droplets at 40% RH when virus was added to PBS alone (IAV Only), or added to PBS containing live wild-type *S. pneumoniae* (+*St.pn*) or live un-encapsulated *S. pneumoniae* (R6), both at 10^8^ CFU/mL final concentration. Droplets were deposited on a non-binding 96-well plate, with triplicate droplets of each mixture recovered at time-points 0, 15, 30, 45, and 60 minutes post-deposition, for quantification and correction as in (A). Individual data points of triplicate droplets are shown, presented as geometric mean ± geometric SD. No significant difference (ns) in infectious titers at time 60 between the two bacteria groups were determined by One-Way ANOVA. **D)** 10 µL of untreated, heat-shocked, or lysed *S. pneumoniae* bacteria at 10^9^ CFU equivalent/mL were stained with crystal violet and deposited on glass slides. Cover slips were fixed on top of wet mounts, and cells were imaged at 100× magnification using an oil immersion objective. Scale bar = 10 µm. Images are representative of duplicate samples. Inset (white box) is zoomed in to highlight individual cells. **E)** Decay of IAV in 1-µL droplets at 40% RH when virus was added to PBS alone (IAV Only), or added to PBS containing live *S. pneumoniae* (+*St.pn*, Live), heat-shocked *S. pneumoniae* (+*St.pn*, Heat), or lysed *S. pneumoniae* (+*St.pn*, Lysed) at 10^8^ CFU equivalent/mL final concentration. Droplets were deposited on a non-binding 96-well plate, with triplicate droplets of each mixture recovered at time-points 0, 15, 30, 45, and 60 minutes post-deposition, for quantification and correction as in (A). Clear symbols indicate samples that were below plaque assay limit of quantification (LOQ), and were set a LOQ/√2. Individual data points of triplicate droplets from 2 independent experiments are shown (n = 6 droplets total per group), with geometric mean ± geometric SD. Significant differences in infectious titers at time 60 relative to IAV only control were determined by Kruskal-Wallis test (*, P ≤ 0.05, ns, not significant). **F)** 1-µL droplets were prepared as in E), then filmed during drying at 40% RH. Images of drying droplets were taken every 16 seconds beginning 2 minutes post-deposition, and diameters of all droplets from each image were quantified by ImageJ processing. Data is presented as the mean diameter (thick line) ± S.D. (shaded error regions) of triplicate droplets, from 2 independent experiments (n = 6 droplets total per group).

Additionally, non-respiratory bacteria *Escherichia coli* (enteric bacterium) and *Pseudomonas syringae* (plant bacterium) were tested for potential stabilisation of IAV in droplets at 40% RH, though data indicated that these alternative bacteria were not significantly stabilising at the tested conditions compared to virus alone (**Supplementary Figure S6**). This suggested that not only did bacteria possess unique qualities relating to virus stabilisation, but it may be that respiratory bacteria in particular were most effective when paired with IAV.

To investigate further, a singular protective respiratory bacterium, *S. pneumoniae*, was chosen for subsequent analyses. Initially, the wild-type encapsulated isolate of *S. pneumoniae* was compared to a genetically modified non-encapsulated mutant. This non-encapsulated mutant (termed R6) is of the same serotype and expresses a similar protein profile to wild-type *S. pneumoniae*, but lacks the *cps* locus required to produce capsular polysaccharide (see **Supplementary Figure S7** for SEM images of each). **Figure 5C** shows that removal of the capsule had no effect on the ability of the bacteria to provide stabilisation for IAV.

Next, *S. pneumoniae* was treated via two physical inactivation methods. One treatment (heat shock) removed cellular viability but retained intact cellular morphology, whilst the other (lysis by prolonged heat-treatment) removed both viability and cellular intactness. Both methods were confirmed to sterilise the bacterial suspensions by CFU plating, and the effect on morphology was confirmed by oil-immersion microscopy (**Fig. 5D**). Inactivated *S. pneumoniae* at 10^8^ CFU/mL equivalent of inactivated bacteria were then compared to the live version for stabilisation of IAV in droplets. **Figure 5E** demonstrates that whilst heat-shock had no detrimental effect – and thus bacterial viability must not be required for enhanced viral stability, bacterial lysis did completely abrogate protective ability. Diameters of drying droplets were also measured at the same 40% RH conditions, with diameters recorded every 16 seconds. Data in **Figure 5F** shows live and heat-shocked *S. pneumoniae* had the same effect on the droplet diameter, retaining a wider droplet diameter compared to droplets of IAV alone.

Conversely, despite containing comparable organic matter, droplets containing IAV plus lysed *S. pneumoniae* showed minimal changes in droplet behaviour relative to IAV-only droplets, with a progressive shrinking in 2D diameter until efflorescence at approximately 20 minutes post-deposition. Collectively, these data show that lysed bacteria were unable to alter droplet drying morphology and were unable to confer stabilisation of the virus.

### Bacterial-mediated virus stabilisation is maintained in a physiologically relevant matrix

An increasing number of studies are emerging that demonstrate the pronounced effect of matrix composition on inactivation kinetics of studied viruses. To ensure the trends we were observing here were able to be replicated in a more physiological matrix, artificial saliva was tested. **Figure 6A** shows a similar inactivation kinetic for IAV alone in artificial saliva droplets compared to the previously tested PBS droplets. Importantly, viral stabilisation by bacteria (*S. pneumoniae*) was also observed in this physiological matrix, despite the increased composition complexity. The protective effect was slightly dampened relative to previous figures where PBS was used, but by the end of the time-course there remained >1.5-log10 more infectious virus in bacteria-containing droplets compared to virus alone. Earlier efflorescence of droplets was also observed in the bacteria case (**Fig. 6B**).

**Figure 6.**
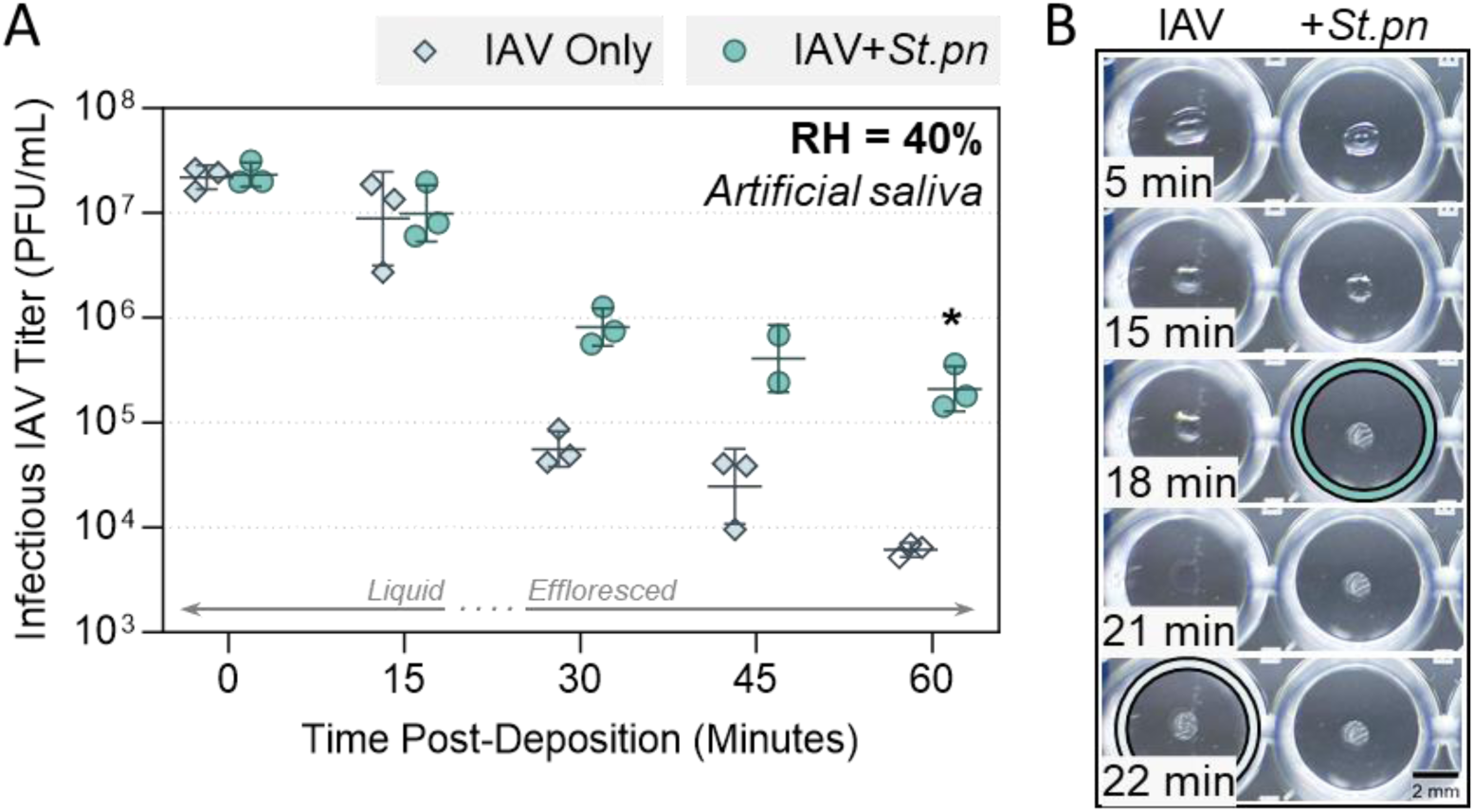
Decay of IAV in 1-μL artificial saliva droplets at 40% relative humidity (RH) and 22 – 25°C in a humidity controlled chamber. **A)** Virus was added to artificial saliva alone (IAV Only), or added to artificial saliva containing live *S. pneumoniae* (IAV+*St.pn*), at 10^8^ CFU/mL final concentration. In both cases, virus was added for 5 × 10^7^ PFU/mL final concentration. 1-μL droplets of each virus suspension were deposited on a non-binding 96-well plate and exposed to indoor air conditions for a total of 60 minutes. Triplicate droplets of each mixture were recovered at time-points 0, 15, 30, 45, and 60 minutes post-deposition, and infectious viral titers were quantified by plaque assay (clear symbols indicate samples that were below plaque assay limit of quantification (LOQ), and were set a LOQ/√2). Infectious viral titers were corrected for physical recovery (determined by genome quantification of IAV in each recovered droplet by dPCR) relative to samples collected immediately after deposition (time 0, where no physical loss has occurred due to drying). Individual data points of triplicate droplets are shown, presented as geometric mean ± geometric SD. Significant differences between IAV titers at time 60 was determined by Mann-Whitney test (*, P ≤ 0.05). **B)** 1 µL droplets containing IAV alone or IAV mixed with *S. pneumoniae* in artificial saliva [as in (A)] were recorded during drying at 40% RH. Images are representative of 3 individual droplets per group. Coloured circles highlight the time-point where crystallisation was first visible by eye for each sample. Scale bar = 2 millimetres (mm).

### Respiratory bacteria are found in the same size fractions as IAV in aerosols, and can mediate partial viral stabilisation

To test whether respiratory bacteria could also be protective within the smaller micro-environment of an expiratory aerosol, suspensions of virus alone or virus mixed with the protective bacterium *S. pneumoniae* were aerosolized and held suspended within a 1.6-m^3^ aerosol chamber at 40% RH. The system used for viral and bacterial aerosolization was designed to mimic the physiological mechanism of bronchiole fluid film burst, or bubble burst, for aerosol generation. This occurs in the lower respiratory tract when the small airways open and close with tidal breathing. *S. pneumoniae* was again selected as the representative bacterium for these experiments as live *S. pneumoniae* has been detected in the lung of healthy volunteers (8–10), where this passive mode of airborne particle generation occurs.

Initially, each inoculum (IAV alone or IAV pre-mixed with *S. pneumoniae,* both in PBS) was aerosolized into the aerosol chamber for 30 seconds, to generate a suitably large aerosol plume. Then, air was collected for a period of 15 minutes into an Andersen multistage Impactor, which collects and fractionates aerosol particles based on their aerodynamic cut off diameter from 0.65 – 7 μm. Each of the 6 size fractions were titrated by plaque assay, and were extracted and quantified for viral genome copies by dPCR. **Figure 7A** shows that for both inoculum types, infectious IAV could be recovered from all 6 of the aerosol size fractions. However when IAV was co-aerosolized with bacteria, three aerosol size fractions between 0.65 – 3.33 μm contained 1-log10 more infectious virus compared with IAV aerosolized alone; these three size fractions are also where the most infectious bacteria were detected. This appeared to be a true increase in infectivity rather than total virions, as comparable numbers of IAV genomes were quantified in these fractions between IAV only and IAV+bacteria experiments. Bacterial genome copies relative to infectious units appear in **Supplementary Figure S8,** with genome copies also slightly enriched in the smaller size fractions.

**Figure 7.**
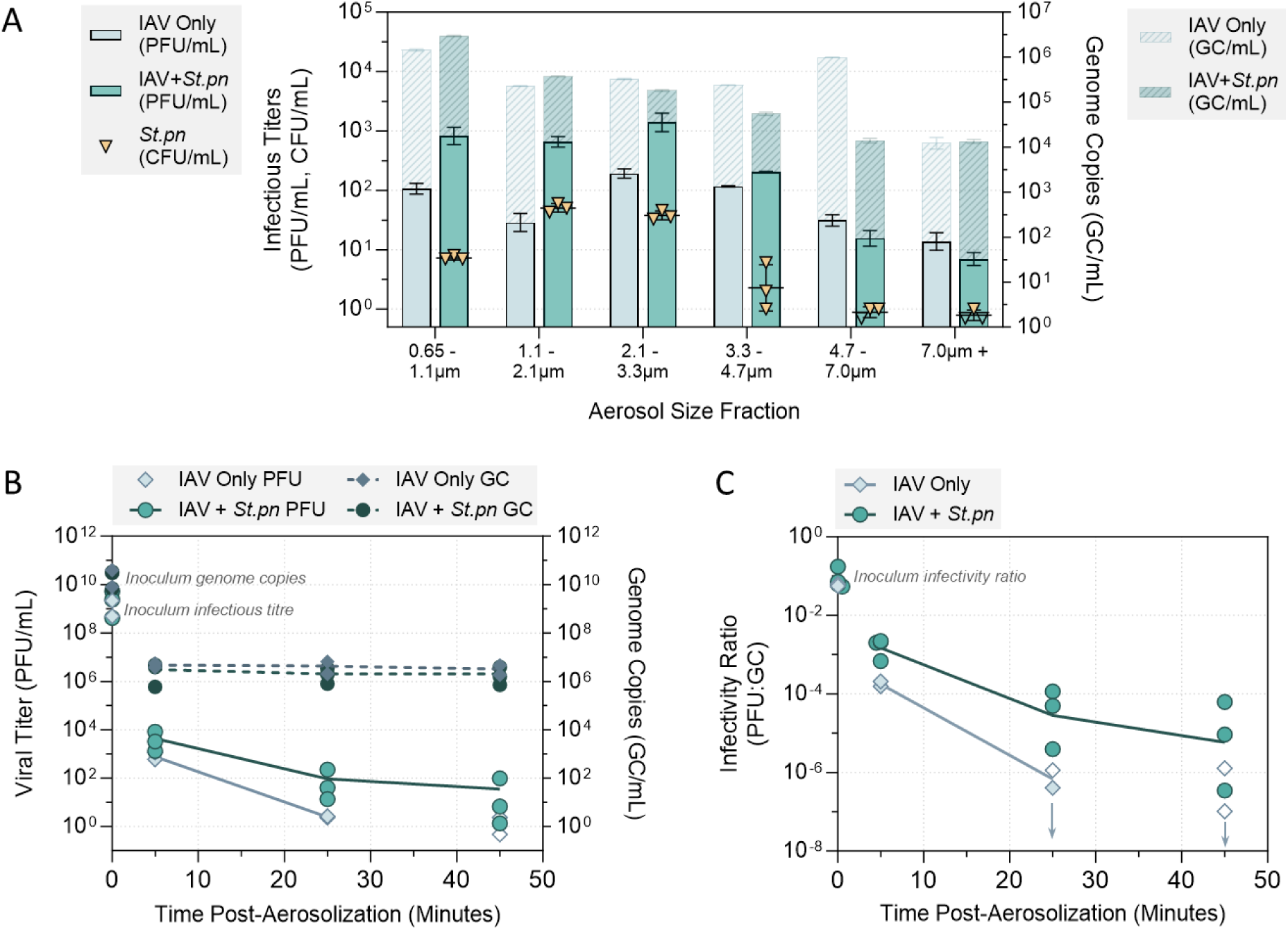
Infectivity of IAV in suspended aerosol particles at 40% RH. IAV was added to PBS alone (IAV Only), or added to PBS containing live *S. pneumoniae* bacteria at 5 × 10^8^ CFU/mL (IAV+*St.pn*). Virus was spiked in at 2 × 10^9^ PFU/mL final viral concentration in both cases. For each experiment, the liquid inoculum (either IAV alone or IAV+*St.pn*) was added to a sparging liquid aerosol generator (SLAG), and nebulised into an aerosol chamber (comprised of a sealed 1.6-m^3^ polytetrafluoroethylene (PTFE) chamber suspended inside a large biosafety cabinet) for a total of 30 seconds, with the air-flow set at 30L air/min. The chamber was maintained at 24 ± 1°C and at the targeted RH of 40% (± 3%) for the full duration of each experiment. **A)** Immediately after the 30-second nebulisation, aerosol particles were recovered using an Andersen Impactor for a total of 15 minutes. Aerosol samples collected by each Stage of the Andersen impactor (6 stages total) were recovered for quantification of infectious IAV (PFU/mL, left axis) and viral genome copies (GC/mL, right axis). Infectious titers were determined in technical triplicate by plaque assay, and viral genome copies were determined in technical duplicate by dPCR. Infectious bacteria (determined by agar plating in technical triplicate) were also recovered from each Stage in the IAV+*St.*pn case, and bacterial titers are overlaid in yellow triangles (CFU/mL, left axis). Clear symbols indicate samples that were below plating limits of quantification (LOQ), and were set a LOQ/√2. Geometric mean ± geometric SD is shown in all cases, with one independent experiment conducted for each inoculum. **B)** IAV only and IAV+*St.pn* inoculums were prepared and nebulised into the aerosol chamber as in (A), but aerosol particles were instead recovered using the Bio-Spot VIVAS. Aerosol samples were collected into the Bio-Spot VIVAS for 10 consecutive minutes, either immediately after nebulisation (collection from 0 – 10 min), 20 minutes after nebulisation (collection from 20 – 30 min), or 40 minutes after nebulisation (collection from 40 – 50 min). Data points are plotted at the mid-point of each collection period (i.e. 5, 25, 45 minutes). For a single experiment, infectious viral titers and viral genome copies were quantified at each time-point by plaque assay and dPCR, respectively. The IAV only inoculum was tested in 2 independent experiments, whilst the IAV+*St.pn* inoculum was tested in three independent experiments. For each independent experimental repeat, infectious virus was quantified in technical triplicate, and genome copies were quantified in technical duplicate. Data points indicate the mean infectious titer (PFU/mL) and mean genome copies (GC/mL) at each time-point from each experiment (clear symbols indicate samples that were below plaque assay limit of quantification (LOQ), and were set a LOQ/√2). Means across independent experiments are also indicated for infectious virus (solid lines) and genome copies (dashed lines). Lines are not drawn when all samples within a group were below LOQ. **C)** Values from experiments in (B) were also used to calculate infectivity ratios. At each time-point of each independent repeat, the mean infectious viral titer (PFU/mL) was divided by the mean recovered virus (i.e. genome copy number, GC/mL) from the matched sample to generate an infectivity ratio. The average infectivity ratio for inoculum virus prior to nebulisation was 8 × 10^-2^ (i.e. 8 infectious particles for every 100 collected particles) across all experimental repeats. Where required, points are nudged horizontally for visibility. Data points indicate infectivity ratios for each independent repeat at each time-point, with means across experiments joined by solid lines for each inoculum type. Clear symbols and downward arrows indicate samples that were below plaque assay LOQ in (B), and connecting lines are not drawn when all samples within a group were below LOQ at a given time-point.

As bacteria and infectious virus were found to be co-localised within the same aerosol fractions by the Andersen sampling method, longer exposure experiments were then performed to assess if aerosolized virus could also be stabilised over longer time-periods by co-localised bacteria. Again, liquid inoculum samples were aerosolized for 30 seconds into the 1.6-m^3^ aerosol chamber, and air samples were now collected into the BioSpot VIVAS instrument for 10 consecutive minutes at 3 individual time-points (0-10 mins, 20-30 mins, 40-50 mins post-aerosolization) for determination of residual viral titers by plaque assay. Samples were also extracted and quantified for viral genome copies by dPCR, as a measure of physical recovery.

**Figure 7B** shows that the total viral genome copies collected did not drop over the course of these 50-min experiments. There was a slight trend for fewer genome copies to be recovered from the aerosolized IAV+bacteria samples, though this difference was within a factor of 2 across matched experiments. Additionally, infectious viral titers were slightly higher when the virus was co-aerosolized with bacteria (**Fig. 7B**). This difference was between 0.5 – 1 log10 at time-points 5 and 25 mins, with a similar if not larger difference at 45 mins (IAV alone was below quantification limits by this time-point in both independent experimental repeats). To account for any potential fluctuations in the total number of virions aerosolized when bacteria were present, infectious titers were also divided by total genome copies recovered in each corresponding sample. This provided a measure of infectious virus relative to total virions recovered (i.e., an ‘infectivity ratio’), and the change in this ratio with time is presented in **Figure 7C**. Data show the infectivity ratio was comparable prior to aerosolization between IAV only and IAV+bacteria inoculums, but post-aerosolization, the presence of bacteria caused ∼0.5-log10 more viral infectivity to be retained at the first sampled time-point. The average infectivity difference was then increased to ∼1.5-log10 by 25 minutes. By 45 minutes, infectivity for IAV aerosolized alone was below detection limits, thus the degree of difference was harder to quantify. Still, IAV aerosolized with bacteria was above the LOQ in all samples by 45 minutes, and thus, these bacteria were also stabilising to IAV even in small aerosol particle.

## Discussion

Airborne transmission of respiratory viruses is a major challenge to address in modern society, and improved understanding of factors that influence the stability of expelled viruses is crucial to mitigate ongoing risks of seasonal epidemics and future pandemics. In the case of both respiratory viruses and enteric viruses, there is a highly complex commensal microbiota also present at the sites of infection. Prior studies have found that commensal gut bacteria can be stabilising to enteric viruses against environmental stresses (14–16), though similar investigation was lacking for resident pathogens of the respiratory niche.

In the present study, experiments primarily focused on 1-µL static droplets that were deposited and exposed to indoor air conditions. Droplets containing IAV mixed with a representative respiratory microbiota (**Fig. 1A**), individual Gram-positive respiratory bacteria (**Fig. 2A**), or certain Gram-negative respiratory bacteria (**Fig. 3A**), demonstrated significant viral stabilisation when compared to virus deposited alone, with 10-100 fold more infectious virus after a full hour of droplet exposure to 40% RH room air conditions. Past work has shown that respiratory commensal bacteria, namely *S. pneumoniae* and *M. catarrhalis*, could be stabilising to IAV in bulk solutions subject to desiccation treatment (13), and our data now show that this stabilisation also occurs in droplet micro-environments.

In the droplet system, one mode of action for viral stabilisation appeared to be the induction of early efflorescence by the large bacteria. Previous work (19) has demonstrated that exposure to high salt molality is the critical driver of IAV inactivation in droplets. Supersaturated salt molalities occur as the droplet progressively evaporates, and rapid virus inactivation occurs under these supersaturation conditions. Efflorescence then reduces the molality of drying droplets back to saturation levels, where inactivation kinetics are comparatively much slower (19). In the current study, droplets containing bacteria effloresced more rapidly compared to droplets containing IAV only (**Fig 2B** and **3B**). This allows the bacteria-containing droplets to pass more rapidly through the supersaturation phase into efflorescence, and the co-localised viruses are quickly removed from an inactivating supersaturated salt environment. In contrast, virus-only droplets progressively move through saturation into supersaturation, and finally effloresce, leaving substantially more time for salt-mediated virus inactivation. Niazi *et al.* (20–22) also found that rapid evaporation and thus rapid efflorescence was stabilising to IAV, human rhinovirus, and respiratory syncytial virus (RSV) in aerosol particles. One can thus speculate that similar bacterially-mediated stabilising effects would also be observed for other respiratory viruses.

Our data show that this early efflorescence was due to altered droplet morphology, rather than a shift in efflorescence relative humidity (ERH) by the presence of bacteria. **Figure 4A** confirmed that bacteria-containing droplets exposed to a rapid decrease in RH effloresced approximately 3 minutes earlier than equivalent droplets without bacteria. However, **Figure 4B** and **4C** show that in slow drying experiments, efflorescence RH between the two droplet types was the same, indicating that the presence of bacteria did not alter ERH. Instead, droplet morphology was consistently observed to be altered by bacterial presence in both fast (**Supp. Videos S1, S2**) and slow drying scenarios (**Supp. Videos S3, S4**). Specifically, bacteria-containing droplets retained larger 2D diameters during the entire drying course, up to the point of efflorescence. This was observed visually for fast drying experiments in **Figures 2B** and **3B**, and quantified in **Supplementary Figure S3** (fast drying) and **Figure 4D** (slow drying). As hygroscopic growth for droplets with and without bacteria should be essentially the same, given by the water activity of PBS, these observed differences in droplet radii were instead due to different droplet morphologies. PBS or virus-only droplets exhibited a shape closer to a half sphere, while the droplets containing bacteria flattened substantially upon drying, with a much smaller change in 2D equivalent radius over time. Flattening of the droplets, and a subsequent increase in surface area to volume ratio, resulted in faster water loss due to evaporation and therefore in earlier efflorescence relative to standard half-sphere drying under identical environmental conditions.

Bacterial lysis experiments revealed that pure organic content was not sufficient to induce this droplet flattening, but rather whole intact bacteria were required for viral stabilisation via the early efflorescence mechanism (**Fig. 5D-F**). This is somewhat similar to observations by Rowe *et al.,* where lysis of bacteria also eliminated bacterial-mediated stabilisation of IAV (13). However, their experiments used bulk solutions and thus are independent of droplet morphology, indicating presence of an additional stabilisation mechanism that is similarly disrupted by bacterial lysis. Interestingly, this prior study also showed that removal of the capsule layer from *S. pneumoniae* was associated with loss of IAV stabilisation, however our data showed no requirement for the capsule (**Fig. 5C**). Differences in micro-environment (droplet vs. bulk solution) and the dominant protection mechanism in each is a likely cause for this discrepancy. Polystyrene spheres of similar shape and size to bacteria also did not cause droplet flattening nor induce early efflorescence here, and when compared to droplets of virus alone, these spheres did not provide any increase in viral stability (**Fig. 5A-B**). Thus, the intact and organic bacteria must possess specific properties that are key to altered drying behaviour of droplets and the associated viral stabilisation.

A similar influence of droplet drying behaviour on IAV stability was observed by Rockey *et al.* (23). Authors tested IAV stability in droplets of human saliva and airway surface liquid (ASL), and observed pronounced decay in saliva at 50% RH (approximately 3-log10 after 2 hours, similar to the total decay observed here in **Fig. 6A** under comparable conditions with artificial saliva), but found relative viral stability in ASL. Authors state this difference in virus inactivation was not associated with bulk protein content nor salt content differences between the two matrices, but instead could be related to the varying droplet behaviours observed in efflorescing conditions. They noted a small difference in efflorescence time between each matrix, with a drying time of 28.2 mins for airway surface liquid (protective) and 32.4 mins for saliva (non-protective) at 50% RH. This is analogous to the difference in efflorescence times seen here between bacteria (protective) and virus only (non-protective) conditions (**Fig. 2B, 3B**, and **4A**).

Combined, it appears that drying droplet morphology and resulting efflorescence time are key determinants of the degree of IAV stability in droplets, and much like different respiratory matrices, commensal respiratory bacteria seem to influence these droplet properties in a species-specific manner. The most protective bacteria tested here were Gram-positive *S. pneumoniae* and *S. aureus,* followed by the Gram-negative *M. catarrhalis* (**Fig. 2, Fig. 3**), with these three also retaining the largest 2D droplet diameters during evaporation (**Supp. Fig. S3**). All bacteria in these experiments were used at an equivalent protein density across experiments (see **Supp. Fig. S9** and associated text), and were grown to a comparable early-log phase prior to use. Differences in total organics or cell densities are therefore unlikely contributors to differential viral stabilisation, and other properties (e.g. Gram-status, charge, surface protein profile, secreted metabolites, etc.) must be present to distinguish protective and non-protective species.

Whilst not respiratory-focussed, Cunliffe *et al*. (24) similarly observed that the presence of bacteria in droplets was correlated with a faster evaporation time relative to droplets of saline matrix alone, and in some instances, their study noted statistically significant differences in evaporation time induced by different bacteria. A potential factor contributing to bacterial-specific droplet behaviour is the production of bacterial biosurfactant. Biosurfactant production has been shown for other Streptococcal species for example, and was found to alter surface tension of droplets (25), which could feasibly influence droplet evaporation in a species-specific manner.

In the enteric space, differing levels of bacterial-mediated protection of associated viruses has also been observed (26). For example, Gram-positive bacteria were more effective stabilisers of murine norovirus than Gram-negative bacteria (27) similar to observations here, whilst Gram-negative bacteria and purified lipopolysaccharide (LPS) were more effective stabilisers of picornaviruses (14). In general, these Gram-negatives have a thick layer of LPS covering their outer membrane, with a thin peptidoglycan (PG) layer underneath, whilst Gram-positives have a thick outer layer of PG, with embedded lipoteichoic acid, teichoic acid, bacterial proteins, and no LPS (28). A study of human and avian influenza viruses showed that LPS could bind to the surface of influenza virus and disrupt its morphology, with both LPS and Gram-negative enteric bacteria being associated with reduced viral stability and reduced long-term aquatic persistence of the tested influenza viruses (29). Potential interaction of respiratory Gram-negative LPS with IAV here may be the reason for a slight reduction in protective efficacy compared to the Gram-positive bacterial counterparts, though this requires validation.

It is worth noting that these enteric studies used bulk solutions (i.e. non-efflorescing conditions which never go past salt saturation), thus the stabilisation mechanisms by bacteria are likely to be different. In fact, viral protection in these studies was in part due to direct binding of certain viruses to the bacterial surface. Viral-bacterial binding has also been shown between IAV and certain commensal respiratory bacteria (30,31), and this interaction could be the additional stabilisation mechanism observed in our study. When early efflorescence of bacteria-containing droplets was prevented by use of high RH (75%), the presence of bacteria was still stabilising to IAV against decay (**Fig. 2C, Fig. 3C**).

In addition to static droplets, it was important to also address aerosols within this study, to test how wide-spread the potential for bacterial stabilisation of expelled IAV could be. **Figure 7A** showed that not only could infectious IAV and infectious *S. pneumoniae* be effectively recovered from an aerosolized plume, but both virus and bacteria seemed to be localised within the same aerosol size fractions. Specifically, airborne particles between 0.65 – 3.3 μm dry size were enriched for both pathogens when they were co-aerosolized, though it is possible that particles outside of the tested size range (i.e., < 0.65 μm) could also contain high levels of IAV. An animal study previously identified both *S. pneumoniae* and IAV within the same expelled aerosol fraction from co-infected ferrets (32), and it is promising that the mechanical system utilised in our study could reproduce these animal-based findings. Stabilisation of IAV by co-aerosolized bacteria was also observed in our data, with IAV retaining higher infectivity for at least 45 minutes in indoor air conditions (40% RH) when bacteria were also present in the inoculum (**Fig. 7B-C**). An animal study observed that the presence of the respiratory microbiota within the nasal cavity of IAV-infected ferrets was crucial for aerosol spread of the virus to non-infected animals. Depleting these animals of their microbiota completely eliminated aerosol-borne spread of IAV, and re-colonising the depleted ferrets with *S. pneumoniae* alone was sufficient to restore aerosol transmission to non-treated animals (13). Whilst viral decay was not impeded as dramatically in aerosol particles as was seen in larger droplets here (likely due to the presence of fewer bacteria), the infectious dose of IAV required for aerosol infection of human volunteers can be as low as 3 TCID50 total (33). Thus, the slight increase in longevity of aerosolized IAV in the presence of bacteria may be sufficient to have a real impact on the number of infectious doses able to spread to new hosts within an indoor space.

In these aerosols, the particles remained airborne for the entire monitoring period, thus bacterial-mediated alteration of droplet morphology when drying on a surface was no longer a factor. However, the bacteria may occupy a significant part of the total volume in these airborne particles, resulting in less water per particle compared to aerosols that contain only viruses. Bacteria-containing aerosols could therefore have a larger surface area to water volume ratio, resulting in faster evaporation and earlier removal of co-localised viruses from a supersaturated salt environment, similar to what was observed for drying droplets. Alternatively, the additional stabilisation mechanism that functioned in the 75% RH droplet case may also dominate in aerosols. Unravelling the precise mechanism(s) of action of bacteria in each microenvironment will be the focus of future work. Bacterial longevity in aerosols has also been investigated in some recent studies (34,35), and this element would be additionally interesting to incorporate for the respiratory bacteria tested here.

We acknowledge that this study comes with a number of shortcomings. As viral-bacterial stabilisation had not been thoroughly investigated for respiratory pathogens, we chose to work with the simplest system possible. This primarily influenced our matrix choice; a basic saline matrix was used to focus on the potential effect of bacteria as an isolated organic component, and to improve reproducibility compared with more complex respiratory solutions. Respiratory matrices are often heterogeneous, thus replicate droplets can have very different physical properties and internal chemistry, such as pH changes through the loss of CO2 or uptake of acidic trace gases from the ambient air (36,37), which could impede identification of a clear bacterial effect. We did test a more physiological matrix of artificial saliva here, and observed pronounced decay of IAV alone (similar to PBS) plus similar levels of viral protection (**Fig. 6A**) and early droplet efflorescence (**Fig. 6B**) mediated by *S. pneumoniae.* An unrelated study investigating *S. pneumoniae* and IAV in small droplets of respiratory matrix conversely found that bacterial presence did not improve viral stability (32). However, this study used complex airway surface liquid collected from cultures of human lung bronchial epithelial cells, and in this fluid, IAV alone was suitably stable already (authors measured 1-log10 decay total over multiple hours). A different study by the same group showed a direct comparison of ASL to saliva, and again found that IAV in ASL droplets was stable, whilst in saliva droplets there was pronounced decay (23), similar to our observations here. Matrix choice is therefore a critical factor, and is the reason for these differing results of bacterial stabilisation between our studies. For IAV, the generation site of infectious particles (e.g. lower airways vs. the nose) is yet to be conclusively identified, so the most representative matrix is not known. Thus, upper respiratory matrices like saliva plus lower respiratory matrices like ASL should both be considered in the field. Results in nasal mucus or a mixture of mucus and saliva would be additionally interesting to obtain, and future work will aim to characterise whether factors present in these more complex fluids can synergistically enhance the observed protective effect of bacteria.

The pathogen concentrations used for studies such as these are also classically difficult to select. There is a trade-off between using concentrations high enough for accurate enumeration from difficult samples, but that also realistically represent what might be carried by human patients. We acknowledge the doses of virus and bacteria used here are high, and likely are higher than those found in a real-life setting. However, enumerating infectious doses from people is difficult, and as a result there is a large range that physiological concentrations are estimated to sit within. A study enumerating numbers of beta-haemolytic streptococci expelled by various respiratory activities found that nose blowing, over coughing or sneezing, released the highest bacterial numbers from human patients. In these experiments, nose blowing resulted in an average expulsion of 1.1 × 10^7^ CFU and a maximum of >1 × 10^9^ CFU (recovered from a sneezed-on handkerchief in 50 mL of media) (38). The concentrations of bacteria used in our experiments (1 × 10^8^ CFU/mL, giving 1 × 10^5^ CFU per deposited droplet) are not so far from these measured bacterial loads. In terms of viral loads, IAV was used at an average of 10^9^ genome copies/mL. Infectious doses from respiratory fluids and swabs from SARS-CoV-2-infected individuals range from 10^2^ to >10^10^ genome copies/mL, depending on the study, participant, and sample type (39–41), and Influenza A and B viral loads have both been detected in clinical specimens across a similar range (41). Future work will investigate whether the trends observed here are retained when the doses of infectious material per particle are reduced, though this will require higher sensitivity enumeration methods. Additionally, this work predominantly utilised the lab-adapted A/WSN/33 strain of IAV. Comparison to the clinically isolated A/Netherlands/07/2009 showed similar inactivation kinetics and similar stabilisation by bacteria (**Supp. Fig. S4**), though it would be beneficial to comprehensively characterise the range of influenza viruses and other respiratory viruses able to be influenced by bacterial presence in droplets and aerosols.

In summary, the COVID-19 pandemic highlighted that many unknowns remain as to how respiratory pathogens effectively spread between hosts. Viruses are vulnerable in the environment, and improved understanding of mechanisms allowing viral persistence will enable us to target this critical transmission window. The composition of the respiratory microbiota may be a previously unconsidered contributing factor towards efficacy of respiratory virus transmission. Identifying the ‘profile’ of bacteria most conducive to emission of stabilised virus could be the focus of future work, may aid in identifying risks of super-spreaders, and ultimately maybe even rationalise differences between viruses with high and low airborne transmission rates. Data indicated at least two mechanisms of bacterial-mediated stabilisation are at play, with one mechanism identified as early droplet efflorescence due to an altered drying morphology. As the mere presence of certain bacteria within a droplet caused earlier efflorescence and heightened virus stability, our findings are also highly relevant to enteric aerosol particles and droplets, e.g. those produced from wastewater or toilet facilities, where bacterial burdens are also high. Furthermore, investigation of expelled pathogen stability using microbially-complex solutions is not widely performed in the current scientific landscape, and should be adopted in the future for both respiratory and enteric-focussed studies to help elucidate the complexity and variability of pathogen transmissions seen in the human population.

## Materials & Methods

### Cells

Madin-Darby Canine Kidney (MDCK) cells (lab of Ben Hale (UZH)) were grown at 37°C in a humidified atmosphere enriched with 5% CO2. Cells were cultured in Dulbecco’s modified Eagle’s medium (DMEM, Gibco), supplemented with 10% heat-inactivated foetal bovine serum (FBS, Gibco), and 100 U/mL of penicillin-streptomycin (P/S, Gibco).

### Virus Stocks

Influenza A virus (IAV) strains A/WSN/33 (lab-adapted H1N1) and A/Netherlands/2009 (clinical H1N1) were inoculated onto confluent MDCK monolayers at a MOI of 0.001. Cells were infected for 72 hours in OptiMEM (Gibco) supplemented with 1% P/S and 1 μg/mL TPCK trypsin (Sigma, Cat. No. T1426). Infected culture supernatants were clarified by centrifugation at 2,500 ×*g* for 10 min, and IAV was then pelleted through a 30% sucrose cushion at 112,400 ×*g* in a SW31Ti rotor (Beckman) for 90 min at 4°C. Pellets were recovered in phosphate buffered saline (PBS, ThermoFisher, Cat. No. 18912014) overnight at 4°C. Concentrated IAV stocks were quantified at 8.0 × 10^10^ and 2.0 × 10^10^ plaque forming units (PFU)/mL (A/WSN/33 and A/Neth, respectively) by plaque assay.

### Bacterial Stocks

Gram-negative strains *Moraxella catarrhalis* (Strain Nell, DSM 9143), *Pseudomonas aeruginosa* (Strain CCEB 481, DSM 50071), *Haemophilus influenzae* (Strain 572, serotype B, DSM 11969) were obtained from DSMZ (Germany). Gram-positive strains *Staphylococcus aureus* (Strain NCTC 8325), *Streptococcus pneumoniae* (Strain D39V, serotype 2), and *Streptococcus pneumoniae* R6 (non-encapsulated derivative of D39V, serotype 2) were a kind gift from Professor Jan Wilhelm-Veening (DMF, University of Lausanne, Switzerland). Stocks were streaked onto agar plates overnight, then healthy colonies used to inoculate and grow liquid cultures to mid-log phase (see **Supplementary Table S1** for media and growth conditions). Once at the required density, liquid cultures were frozen in growth media supplemented with 15 – 20% glycerol, and stored at -80°C. When required for experiments, bacterial cells were thawed on ice, washed 3× in PBS to remove glycerol and residual media components (10,000 ×*g*, 4° C, 5 mins), then resuspended in the required matrix (PBS, or artificial saliva) at 1 × 10^8^ colony forming units (CFU)/mL for immediate use (see **Supp. Fig. S9** for the titer correction made for *M. catarrhalis*). Artificial saliva (ASTM E2720-16 composition) was obtained from Pickering Laboratories, USA, Cat. No. 1700-0317 (see **Supplementary Table S2** for composition).

### Enumeration of Microorganisms

Plaque assay for viral titration was conducted using monolayers of MDCK cells in 12-well plates. IAV samples were serially diluted in ‘PBS for infections’ (PBSi; PBS supplemented with 1% P/S, 0.02 mM Mg^2+^, 0.01 mM Ca^2+^, and 0.3% bovine serum albumin [BSA, Sigma-Aldrich, Cat. No. A1595], final pH ∼7.3), before being added to washed cellular monolayers. Viruses were incubated on monolayers for 1 h at 37°C with 5% CO2 with manual agitation every 10 min. Non-attached viruses were removed from cells, and MEM supplemented with 0.5 μg/mL TPCK-trypsin and agarose was added to cells. Infected and control plates were incubated for 72 h at 37°C with 5% CO2, and plaques were visualized after fixing cells in PBS + 10% formaldehyde (Sigma, Cat. No. 47608-1L-F), then staining with 0.2% crystal violet solution (Sigma, Cat. No. HT901-8FOZ) in water + 10% methanol (Fisher Chemical, Cat. No. M-4000-15). Depending on the volume of non-diluted sample plated in each titration assay (between 100 μL to 1.2 mL), the limit of quantification (LOQ) varied between 10 PFU/mL to 0.67 PFU/mL. Positive values were counted only when two or more samples out of a triplicate showed visible plaques. Data points below each corresponding LOQ were set at LOQ/√2, as recommended for non-detectable values by Hornung and Reed (42).

Bacterial stocks and experimental samples were enumerated for viable bacteria by serially diluting in PBS or growth media. Stock samples were titrated by spot-plating 25 μL of each dilution onto agar plates, whilst experimental samples were titrated by spreading 100 μL – 1 mL of solution directly onto agar plates in triplicate. Where experimental samples had low concentrations, samples were plated undiluted. After overnight incubation of plates, colonies were enumerated by eye. See **Supplementary Table S1** for agar types used for each bacterial species. Depending on the volume of non-diluted sample plated (100 μL – 1 mL), the limit of quantification (LOQ) varied between 10 CFU/mL to 1 CFU/mL. Positive values were counted only when two or more samples out of a triplicate showed visible colonies. Samples with no detectable bacteria were set at LOQ/√2 to differentiate between samples at the LOQ and those with no detection, as recommended for non-detectable values by Hornung and Reed (42).

### Viral genome quantification

RNA extraction was performed using the QIAamp Viral RNA Mini extraction kit (Qiagen, Cat. No. 52906) according to manufacturer’s instructions. Viral RNA was stored at -20°C until analysis by digital PCR (dPCR). Amplification and detection were performed using the QIAcuityTM OneStep Advanced Probe Kit (Qiagen, Cat. No. 250132), with primers and probe targeting the influenza A virus M gene segment (specific amplicon length is 110 bp), based on the assay by Ward *et al.* (43). Primers and probe were obtained from Microsynth (Switzerland), and were reconstituted in Ultrapure RNase-free water according to manufacturer’s instructions, to make 100µM stocks (stored stocks at -20°C until use). When required, primers and probe were mixed in RNase-free water to make a combined Primer-Probe working solution (forward and reverse primers at 8 µM, probe at 4 µM final concentration).

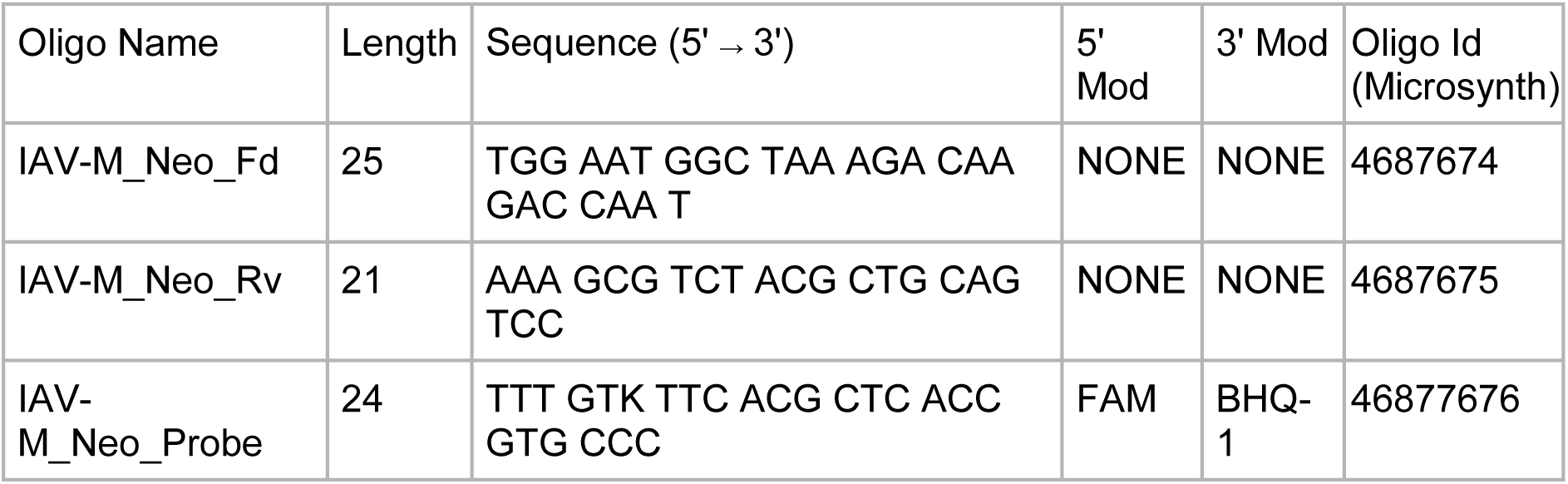

The dPCR mixture for a single reaction for IAV detection was 4 µL of 4x OneStep Advanced Probe Mastermix, 0.12 µL 100× OneStep Advanced RT Mix, 1.2 µL Primer-Probe working solution (giving 0.8µM final concentration of each primer and 0.4 µM final concentration for probe), 1.5 µL Enhancer GC, 2.18 µL RNase-free water, and 3 µL extracted RNA sample. When required, extracted RNA samples were first diluted in RNase-free water to be within dPCR working range (<10^4^ copies/µL to be accurately quantified). Sample dilutions also ensured no PCR inhibition in our experimental protocol. All sample manipulations were performed under biosafety cabinets. Once prepared, the reactions were loaded into QIAcuityTM Nanoplate 8.5k 96-well (Qiagen, Cat. No. 250021) run using the QIAcuity One Digital PCR System (Qiagen, 2-plex Device, Cat. No. 911001). The following conditions were used for the one-step cycling dPCR program: 40 min at 50°C for RT, 2 min at 95°C for enzyme denaturation, followed by 40 cycles of 95°C for 5 sec, then 58°C for 30 sec for annealing and extension. Signal was quantified and copies/uL calculated using the integrated QIAcuity Software Suite. All dPCR runs contained extraction controls and non-template controls which were always negative. For correction of plaque assay data based on physical recovery, PFU data from individual droplets were normalised based on dPCR genome copies recovered at time 0 for each corresponding group, where no physical loss occurs due to adsorption (see (19) illustrating genome copies from a 1 µL deposited droplet that is immediately recovered from the non-binding plate is identical to genome copies quantified when 1 µL is spiked directly into recovery medium).

### Bacterial Genome Quantification

Samples were extracted using the QIAGEN PowerLyzer PowerSoil Kit (Cat. No. 12855-50) according to manufacturer’s instructions. 140 µL of liquid sample was added per tube for extraction, and eluted in 100 µL for storage at -20°C until quantification. Bacterial DNA was quantified using the Femto Bacterial DNA Quantification Kit (Cat. No. E2006) by RT-qPCR and converted from ng to genome copies, all according to manufacturer’s instructions. Each sample was run in technical duplicates, and included a no template control (NTC) and an extraction blank to confirm the absence of contamination. The assay was calibrated over a range of 180 -1.8 × 10^7^ GC/µL, yielding a standard curve with a slope of -3.67, and an intercept of 37.8.

### Droplet Experiments

Experiments in a sealed humidity and temperature controlled chamber (Electro-Tech Systems, Cat. No. 5532) were conducted based on methodology established in (19). IAV stocks were diluted in the appropriate matrix (PBS or artificial saliva ± mucin) to 8 × 10^8^ PFU/mL. Washed bacterial strains were resuspended at 10^8^ CFU/mL in the appropriate matrix, and the diluted IAV was spiked in to give 5 × 10^7^ PFU/mL final viral concentration. Where required, bacteria were mixed together (microbiota case) or inactivated via various treatments (heat-shocked at 95°C for 15 minutes, or lysed by prolonged heat-treatment at 95°C for 30 minutes) prior to addition of the virus. Alternatively, IAV was spiked into matrix alone (no bacteria) at the same final concentration of 5 x 10^7^ PFU/mL. In the case of inorganic polystyrene beads (Cat. No. PP-10-10, 5% w/v, 1.23 µm mean diameter, ∼4.89 × 10^10^ particles/mL, obtained from Spherotech, Illinois USA), beads were diluted to 10^8^ particles/mL in PBS prior to the addition of virus. Virus and bacteria/bead mixtures were kept on ice for ∼10 minutes whilst the humidity chamber equilibrated. Relative humidity (RH, %) and temperature (T, °C) within the sealed humidity chamber were recorded using a portable hygrometer (Model T174H, Cat. No. 05726560, Testo, USA), with readings taken once every minute (confidence intervals provided by the manufacturer were ±0.5 °C, ±3 %RH for T and RH measurements respectively). When required, humidity was adjusted using distilled water, to keep RH levels constant for the duration of the experiment. Once the required RH and temperature were reached, virus solutions were manually mixed immediately prior to deposition of 1 μL droplets on open 96-well plates (non-binding microplates, flat-bottom, transparent, Greiner Bio-One, Cat. No. 7655901, Huberlab, Switzerland). Time 0 (t0) droplets were deposited last within each group, and were immediately recovered from the plate by addition of 300 μL PBSi directly to the wells. Remaining droplets were recovered 15, 30, 45, 60, or 120 minutes after deposition, again by adding 300 μL PBSi to the respective wells. For recovery of all droplets, a 1 mL pipette was used to pipette the 300 μL volume up and down 5 times, followed by physical scratching of the plate surface using the same pipette tip to dislodge any stuck material. This was repeated twice prior to transfer of the full 300 μL to Eppendorf tubes for downstream processing. For each sample, half the volume was frozen for extraction and dPCR, whilst the other half was frozen for titration by plaque assay. As a control, IAV was spiked into the respective matrix and kept in a sealed Eppendorf tube within the humidity chamber, then sampled at the initial and final time-point of the experiment to ensure no viral decay occurred.

When droplet filming was required, a custom 3D-printed stand housing a Raspberry Pi (Raspberry Pi 4 model B Rev. 1.4, with ARMv7 Processor rev 3 v7l) and a mounted Sony camera (Model: IMX477R, 12.3 megapixels, specifications available here: https://www.raspberrypi.com/products/raspberry-pi-high-quality-camera/) were used. The distance from the camera to the non-binding plate surface was 15 cm. Images were acquired every 16 seconds for a total of 30 minutes. Images were then processed using Python CellPose particle identification software (with a custom model pre-trained on droplet images) paired with the particle measurement function in ImageJ.

### Determination of ERH in Droplets

Additional equipment was used to monitor efflorescence of droplets of PBS alone, or PBS containing *S. pneumoniae* bacteria at 10^8^ CFU/mL. For biosafety reasons, bacteria were heat-inactivated (95°C for 15 minutes), then frozen at - 20°C prior to use in experiments. 1 µL droplets of each matrix were manually deposited on a hydrophobic cover slip, or droplets were deposited with a droplet-on-demand printer on a hydrophobic cover slip. Cover slips were then placed in an environmental cell with precision controlled temperature and relative humidity. Full details of the setup are given by Song *et al.* (44). During controlled humidity cycles at constant temperature, morphological changes of the droplets were monitored optically with a microscope (Olympus BX-40, magnification 50× for printed droplets, magnification 4× for 1 µL droplets) equipped with a Raspberry Pi high resolution camera to acquire images and movies of the droplet morphology.

### Aerosol Experiments

Inactivation of IAV in levitated aerosol particles was determined by nebulizing the virus in a 1.6-m^3^ polytetrafluoroethylene (PTFE) chamber suspended inside a large biosafety cabinet. Dry purified air filtered with a HEPA and activated carbon filter was used to flushed the chamber prior to experiments. A scanning mobility particle sizer (SMPS), consisting of a differential mobility analyzer (DMA model TSI long, TSI Inc., Shoreview, MN, USA) and a condensation particle counter (CPC model 3772, TSI Inc., Shoreview, MN, USA), were used to monitor the particle size distribution within the chamber in real-time. A Nafion humidifier (Model FC100-80-6MKS, Permapure LLC, Lakewood, NJ, USA) was used to control the RH in the chamber. Nebulization was performed using a sparging liquid aerosol generator (SLAG, CH Technologies Inc.) and aerosol particles were recovered with either an Andersen Impactor (TE-10-800 Six Stage Impactor System, Tisch Environment, USA), or a viable virus aerosol sampler (BioSpot-VIVAS, Aerosol Devices Inc.). The advantage of the SLAG nebulizer is the formation of aerosol particles by bursting bubbles from a liquid, mimicking the formation of expiratory aerosol particles in the human body. To avoid any leakage, the chamber was never completely inflated when nebulizing infectious viruses. An air ionizer (Model SL-001, Dr. Schneider Holding GmbH, Kronach-Neuses, Bavaria, Germany) was used to balance the formation of electrostatic charges on the chamber wall, in order to limit wall losses of aerosol particles.

In all cases, aerosol experiments were conducted using PBS as the matrix. IAV was added to plain PBS, or added to PBS containing live *S. pneumoniae* bacteria at 5 × 10^8^ CFU/mL (pre-washed 2× in PBS prior to use). Virus was spiked in at 2 × 10^9^ PFU/mL final viral concentration in both cases. For each experiment, 22mL of inoculum was added to the SLAG for nebulisation. Nebulisation into the aerosol chamber was performed for a total of 30 seconds, with the air-flow set at 30L air/min. The chamber was maintained at 24 ± 1°C and at the targeted RH of 40% (± 3%) for the full duration of each experiment. Sampling was performed using either an Andersen Impactor or BioSpot VIVAS, as detailed below. After each experiment, the chamber was decontaminated by UV radiation and ozone-rich air flushing for an hour.

### Size-resolved Aerosol Sampling

For collection of aerosols based on size, the Andersen Impactor (TE-10-800 Six Stage Impactor System, Tisch Environment, USA) was utilised with the aerosol chamber described above. 9 mL of Tryptic soy Broth (Cat. No. 211825, BD) was added to 6 glass petri dishes, with one dish placed underneath each of the 6 Stages during Impactor assembly. The virus alone or virus + bacteria inoculums were nebulised for 30 seconds as described above, then aerosols were collected from the chamber directly into the Andersen Impactor for 15 consecutive minutes immediately after 30 seconds nebulization. The air flow into the Impactor was 28.3 L/min. After collection, the liquid collection media in each layer was transferred to 50 mL conical Falcon Tubes for transport on ice. 7.6 mL of liquid was recovered per stage due to evaporation during 15 minutes of collection. Of this volume, 2.5 mL of sample was titrated immediately by plaque assay (100 μL non-diluted sample plated in each well of 12-well plate, with 2 plates used per Stage) to determine viable IAV concentrations. Where required, samples were also diluted in PBSi to enable enumeration of countable plaques. 3.5 mL was utilised for bacterial agar plating, with 1 mL of non-diluted sample spread and adsorbed onto triplicate agar plates (3× 1 mL per agar plate for each Stage) to enable a low detection limit. Where required, samples were diluted in tryptic soy broth and 100 μL plated to allow for countable colonies. Additionally, 140 μL of sample was frozen for viral RNA extraction and dPCR, and 140 μL of sample was frozen for bacterial DNA extraction and bacterial DNA quantification. Both genome quantification methods are described above.

### Total Aerosol Sampling

For collection of aerosols across a time-course, the viable virus aerosol sampler (BioSpot-VIVAS, Aerosol Devices Inc.) was utilised with the aerosol chamber described above. The virus alone or virus+bacteria inoculums were nebulised for 30 seconds as described above, then aerosols were collected from the chamber directly into VIVAS for 10 consecutive minutes immediately after 30 seconds nebulization. The air flow into the VIVAS was 8 L/min. Aerosol particles were condensed in the VIVAS sampler and virions were collected in a Petri Dish containing 2.5 mL of PBSi. After collection of this initial time-point (0 – 10 min), the 2.5 mL of collected sample was transferred to a 50 mL Falcon tube and kept on ice until the end of the experiment. Additional time-points of 20 – 30 min and 40 – 50 min were collected by the VIVAS in the same manner. Time-points are annotated in the text as the mid-point of each, i.e. 5, 25, and 45 min. All samples were kept on ice until the end of the experiment, and were then aliquoted and frozen at -20°C until quantification. Infectivity was quantified in technical triplicate by plaque assay, whilst genomic copies were quantified in technical duplicate by dPCR.

### Qubit Protein Measurement

Frozen bacterial stocks were thawed and washed 3× in PBS to remove glycerol and residual media components (10,000 ×*g*, 4°C, 5 mins), then resuspended in miliQ H2O at 10^8^ CFU/mL for immediate use (see **Supp. Fig. S9** for the titer correction made for *M. catarrhalis*). Samples were inactivated at 85°C for 20 minutes, then cooled and measured for total protein by Qubit 4 using the Protein Assay Quantification Kit (Life Technologies, Q33211), according to manufacturer’s instructions.

### Scanning Electron Microscopy (SEM)

Bacterial strains to be imaged were thawed on ice, washed 3× in PBS, and resuspended in filter-sterilised PBS at a concentration of 10^10^ CFU/mL. Samples were drop-cast onto SEM supports and left to adhere for 15 minutes. Samples were fixed in glutaraldehyde (1.25% final concentration, 0.1M phosphate buffer, pH 7.4) for 1-2 hours, then washed 3× in cacodylate buffer (0.1M, pH 7.4) for 2 minutes per wash. Samples were stained with osmium tetroxide in cacodylate buffer (0.1M, pH 7.4) for 30 minutes, then washed 3× in H2O for 2 minutes per wash. Samples were then progressively dehydrated in a graded alcohol series (1× 30%, 1× 50%, 1× 70%, 1× 96%, 2× 100%), with 3 minutes of incubation per change. Critical point dry was then performed, and samples were mounted and coated with gold-palladium. Samples were imaged using the Zeiss Merlin Resolution SEM at the BioEM Facility at EPFL.

### Statistical Analysis

Quantitative results are expressed as geometric mean ± geometric standard deviation. Where data was left-censored due to detection limits being reached, non-parametric tests were utilised. Specifically, Mann-Whitney test (two-tailed) was used for comparison of data from 2 groups, and Kruskal-Wallis test (with Dunn’s statistical hypothesis test for correction of multiple comparisons) was used for analysis of data from 3 or more groups involving a single independent variable, according to recommendation from (45). Where data was not left-centred and thus normally distributed, the parametric paired or unpaired t-test, or One-Way ANOVA (with Holm-Šídák’s multiple comparisons test) were used. All analyses were performed using GraphPad Prism, version 10.0.2. (GraphPad Software, La Jolla, USA). P-values ≤ 0.05 (95% confidence) were considered statistically significant.

## Acknowledgements

This work was funded by the Swiss National Science Foundation (Grant #189939 to TK, and Grant #209808 to SD). We thank Marie Croisier (BioSEM specialist, Biological Electron Microscopy Facility, EPFL) for SEM sample preparation and image acquisition.

## Data Availability Statement

All data relating to this study is shared in accordance with FAIR (findable, accessible, interoperable, and reusable) data principles, and data are deposited in the community-approved and cross-disciplinary public repository Zenodo (https://doi.org/10.5281/zenodo.10613395).

