## Supplementary Information and Figures for "Stability of influenza A virus in droplets and aerosols is heightened by the presence of commensal respiratory bacteria"

**Supplementary Table S1** – List of agar, growth media, and growing conditions used for bacterial strains in this study to generate frozen working stocks. Optical density at 600 nm ($OD_{600}$) used to inoculate liquid cultures (initial) and $OD_{600}$ at the mid-log phase (where liquid cultures were halted) are also indicated, with approximate growth times. All working stocks were stored at -80ºC with glycerol prior to use.

| Bacterium | Agar Type | Liquid Media Type | $OD_{600}$ (initial, mid-log) | Growth Conditions | Time until mid-log | Glycerol for storage |
|---|---|---|---|---|---|---|
| *Pseudomonas aeruginosa* (DSM 50071) | Tryptic Soy Agar (TSA, BD 236950) | Tryptic Soy Broth (TSB, BD 211825) | 0.01, 0.60 | 37°C, 200 rpm orbital shaking | 4 – 5 hours | 20% |
| *Staphylococcus aureus* (NCTC 8325) | TSA or Brain Heart Infusion Agar (BHIA, BD 211065) | TSB or Brain Heart Infusion Broth (BHIB, BD 237500) | 0.05, 0.20 | 37°C | 3 – 4 hours | 16% |
| *Streptococcus pneumoniae* (D39V, and R6) | TSA or BHIA | TSB or BHIB | 0.05, 0.25 | 37°C, 5% $CO_2$ | 3 – 4 hours | 16% |
| *Moraxella catarrhalis* (DSM 9143) | BHIA | BHIB | 0.05, 0.50 | 37°C, 5% $CO_2$, 200 rpm orbital shaking | 4 – 5 hours | 20% |
| *Haemophilus influenzae* (DSM 11969) | BHIA+100mg/L Hemin (Sigma-Aldrich H9039), and 2mg/L β-NAD (Sigma-Aldrich N062) | BHIB+100mg/L Hemin (Sigma-Aldrich H9039), and 2mg/L β-NAD (Sigma-Aldrich N062) | 0.05, 0.40 | 37°C, 5% $CO_2$ | 3 – 4 hours | 16% |

**Supplementary Table S2** – Artificial Saliva composition. Artificial saliva product (ASTM W2720-16, non-stabilised, Cat. No. 1700-0317) and composition information were obtained from Pickering Laboratories, USA.

| Component | Concentration per litre of final product |
|---|---|
| Sodium Chloride | 0.88 g/L |
| 0.2 M Potassium Phosphate Monobasic | 7.7 mL/L |
| 0.2 M Potassium Phosphate Dibasic | 12.3 mL/L |
| Potassium Chloride | 1.04 g/L |
| Potassium Thiocyanate | 0.19 g/L |
| Calcium Chloride Monohydrate | 0.13 g/L |
| Magnesium Chloride Heptahydrate | 0.04 g/L |
| Ammonium Chloride | 0.11 g/L |
| Sodium Bicarbonate | 0.42 g/L |
| Urea | 0.12 g/L |

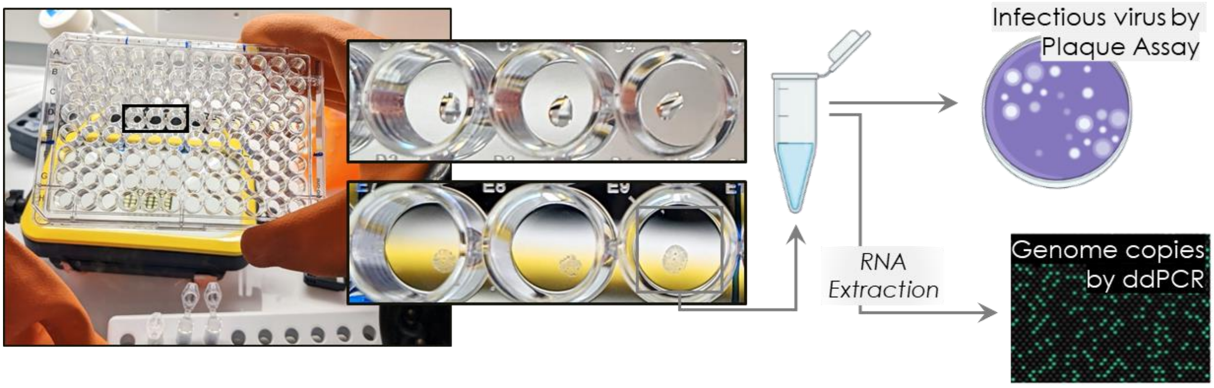


**Supplementary Figure S1** – Standard work-flow of an individual droplet experiment. At time 0, up to 15× 1-µL droplets per group were deposited in individual wells of a hydrophobic 96-well plate. 3× droplets were immediately recovered into 300 µL of PBSi as time 0 controls, and transferred into sealed Eppendorf tubes. The plate was contained within a humidity and temperature controlled chamber during droplet deposition, and for the entire monitoring time-course. As time progressed, droplets would shrink and effloresce (inset) due to equilibration with air. At 15-minute or 60-minute intervals (depending on the RH of interest), these exposed droplets were also collected into PBSi, again with 3 droplets collected at each time-point per group. After collection of all droplets, each recovered sample was split into two aliquots: half of each sample was used for infectivity titration by plaque assay, and the other half was used for RNA extraction and genome copy quantification by ddPCR. Thus, each individual droplet was quantified for both infectivity and physical recovery. All infectivity data was corrected for physical loss relative to time 0 controls. Schematic was made using Biorender.com.

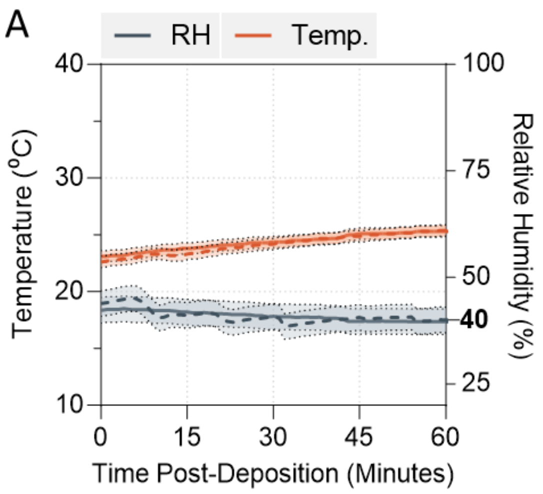


**Supplementary Figure S2** – During all droplet experiments, relative humidity (RH) and temperature (T) were monitored by a portable hygrometer, with readings taken every minute. Representative plots for experiments conducted at either **A)** 40% RH for 60 minutes, or **B)** 75% RH for 120 minutes are shown. Solid and dotted lines distinguish readings from two individual experimental repeats. Confidence intervals of ±0.5 for T (°C), and ±3 for RH (%) (as provided by the manufacturer) are indicated by shaded regions. Bold text indicates the target RH in each case.

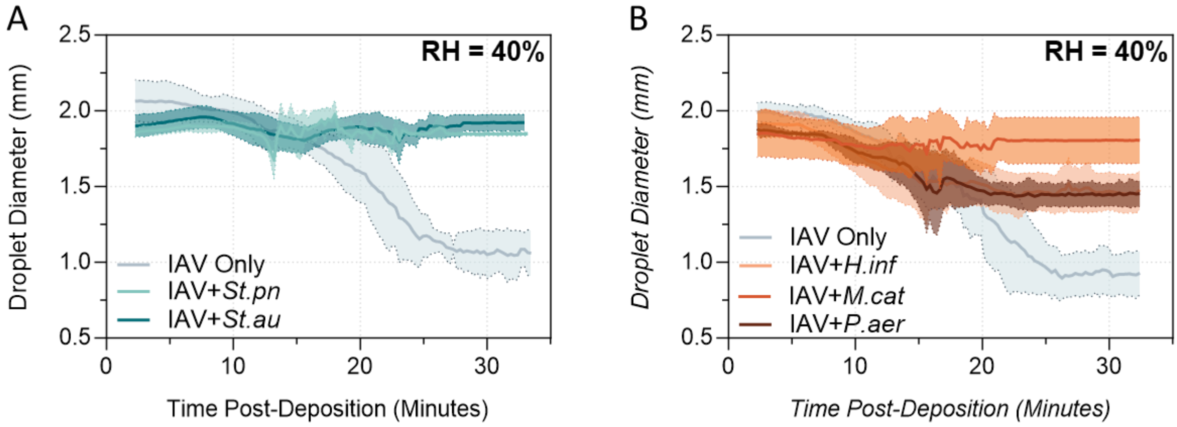


**Supplementary Figure S3** – Diameter measurements of evaporating droplets at 40% RH. **A)** Virus was added to PBS alone (IAV Only), or added to PBS containing live Gram-positive bacteria *Streptococcus pneumoniae* (+*St.pn*) or *Staphylococcus aureus* (+*St.au*) at $10^8$ CFU/mL. In all cases, virus was added for $5 \times 10^7$ PFU/mL final concentration. 1-µL droplets of each virus suspension were deposited on a non-binding 96-well plate and exposed to indoor air conditions for approximately 30 minutes at 40% RH. Droplets were filmed, with images taken once every 16 seconds. ImageJ software was used to extract droplet diameter measurements from each static image. Data shows mean droplet diameter (in millimetres, mm) for each group, with SD indicated by shaded regions (n = 3 or 6 individual droplets recorded per group). Data is combined from 2 independent experimental repeats. **B)** As above, but virus was added to PBS alone (IAV Only), or added to PBS containing live Gram-negative bacteria *Haemophilus influenzae* (+*H.inf*), *Moraxella catarrhalis* (+*M.cat*), or *Pseudomonas aeruginosa* (+*P.aer*) at $10^8$ CFU/mL final concentration.

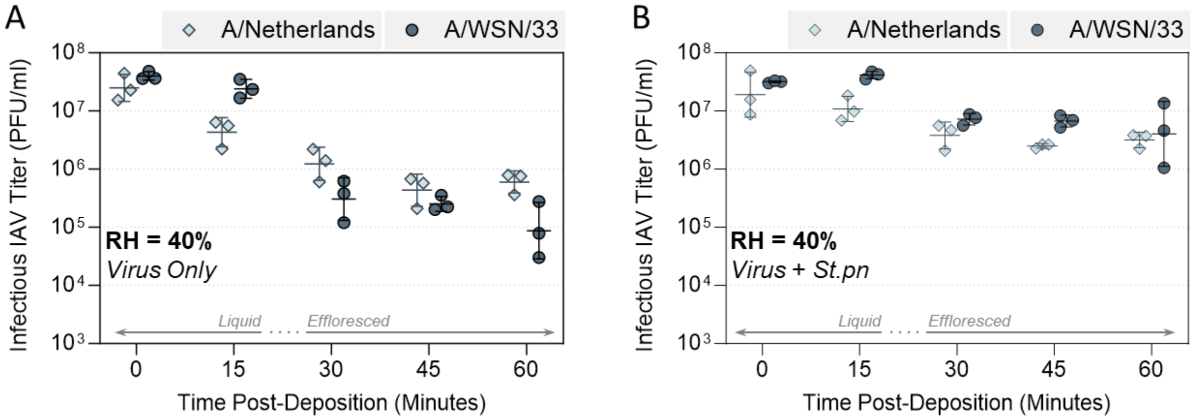


**Supplementary Figure S4** – Comparison of inactivation kinetics of IAV H1N1 strains A/Netherlands/2009 (clinical isolate) and A/WSN/33 (lab-adapted strain) in 1-µL PBS droplets at 22 – 25ºC in a humidity controlled chamber. **A)** Each strain of IAV was added to PBS alone or **B)** added to PBS containing live *Streptococcus pneumoniae* (+*St.pn*) bacteria at $10^8$ CFU/mL. In all cases, virus was added for 5 × $10^7$ PFU/mL final concentration. 1-µL droplets of each virus suspension were deposited on a hydrophobic 96-well plate and exposed to indoor air conditions for a total of 60 minutes at 40% RH. Triplicate droplets of each mixture were recovered at time-points 0, 15, 30, 45, and 60 minutes post-deposition, and infectious viral titers were quantified by plaque assay. Infectious viral titers were corrected for physical recovery (determined by genome quantification for A/Neth and A/WSN/33 in each recovered droplet by ddPCR) relative to samples collected immediately after deposition (time 0, where no physical loss has occurred due to drying). Individual data points of triplicate droplets are presented, with geometric mean ± geometric SD. Significant differences in infectious titers at time 60 relative to IAV only control were tested by One-Way ANOVA, with no significant differences between IAV strains.

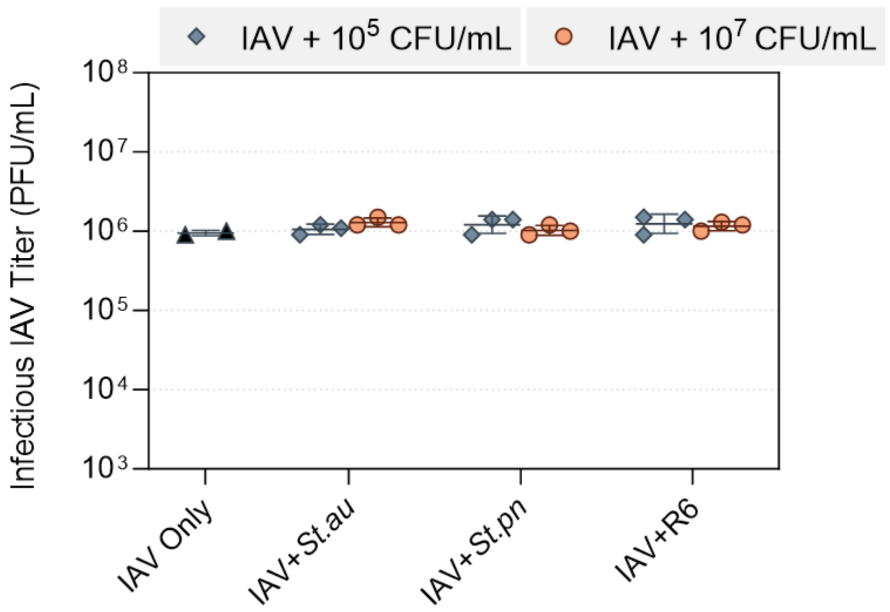


**Supplementary Figure S5 –** Bacterial presence has no direct effect on viral titers by plaque assay. IAV was mixed in bulk solution with *Staphylococcus aureus* (+*St.au*), *Streptococcus pneumoniae* (+*St.pn*), or non-encapsulated *S. pneumoniae* (+R6) bacteria at two different concentrations as specified (equivalent to 1: 0.1 and 1: 10 ratio of virus: bacteria), in PBS. Alternatively, IAV was suspended in PBS alone (IAV Only). These mixtures were then added directly to cell monolayers at a concentration of $1 \times 10^6$ PFU/mL for plaque assay, and after 1-hour adsorption time, bacteria and unbound virus were removed. Antibiotics in the plaque assay media ensured no bacterial growth occurred to affect the health of cells during the subsequent incubation period. Compared to cells infected with IAV alone, the presence of bacteria during the 1-hour infection period at the tested concentrations did not alter infectious viral titers enumerated by plaque assay. Each group was titrated in technical duplicate (IAV only) or triplicate (remaining groups), and data is presented as geometric mean ± geometric SD. Note that in droplet experiments, virus and bacteria are used at a ratio of 1:2 virus:bacteria, which is within the range of concentrations tested here.

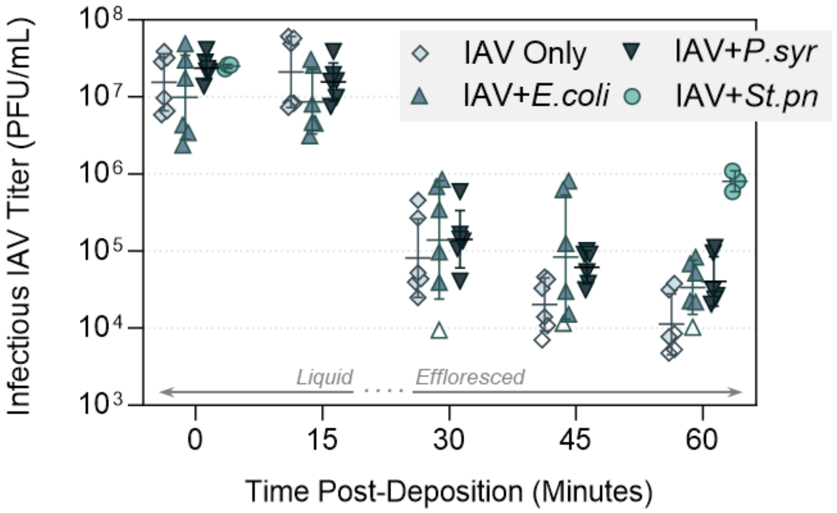


**Supplementary Figure S6** – Decay of IAV in 1-µL PBS droplets at 22 – 25°C in a humidity controlled chamber when mixed with non-respiratory bacteria. Virus was added to PBS alone (IAV Only), or added to PBS containing live *Escherichia coli* (+*E.coli*), *Pseudomonas syringae* (+*P.syr*), or *Streptococcus pneumoniae* (+*St.pn*, positive control) bacteria at $10^8$ CFU/mL final concentration. In all cases, virus was added for $5 \times 10^7$ PFU/mL final concentration. 1-µL droplets of each virus suspension were deposited on a hydrophobic 96-well plate and exposed to indoor air conditions for a total of 60 minutes at 40% RH. Triplicate droplets of each mixture were recovered at time-points 0, 15, 30, 45, and 60 minutes post-deposition (positive control recovered at time 0 and 60 only), and infectious viral titers were quantified by plaque assay (clear symbols indicate samples that were below plaque assay limit of quantification (LOQ), and were set a LOQ/√2). Infectious viral titers were corrected for physical recovery (determined by genome quantification of IAV in each recovered droplet by ddPCR) relative to samples collected immediately after deposition (time 0, where no physical loss has occurred due to drying). Data shows individual data points of triplicate droplets, from 2 independent experimental repeats (n = 6 droplets total per group, with the exception of IAV+*St.pn* which includes just 3 droplets), presented as geometric mean ± geometric SD.

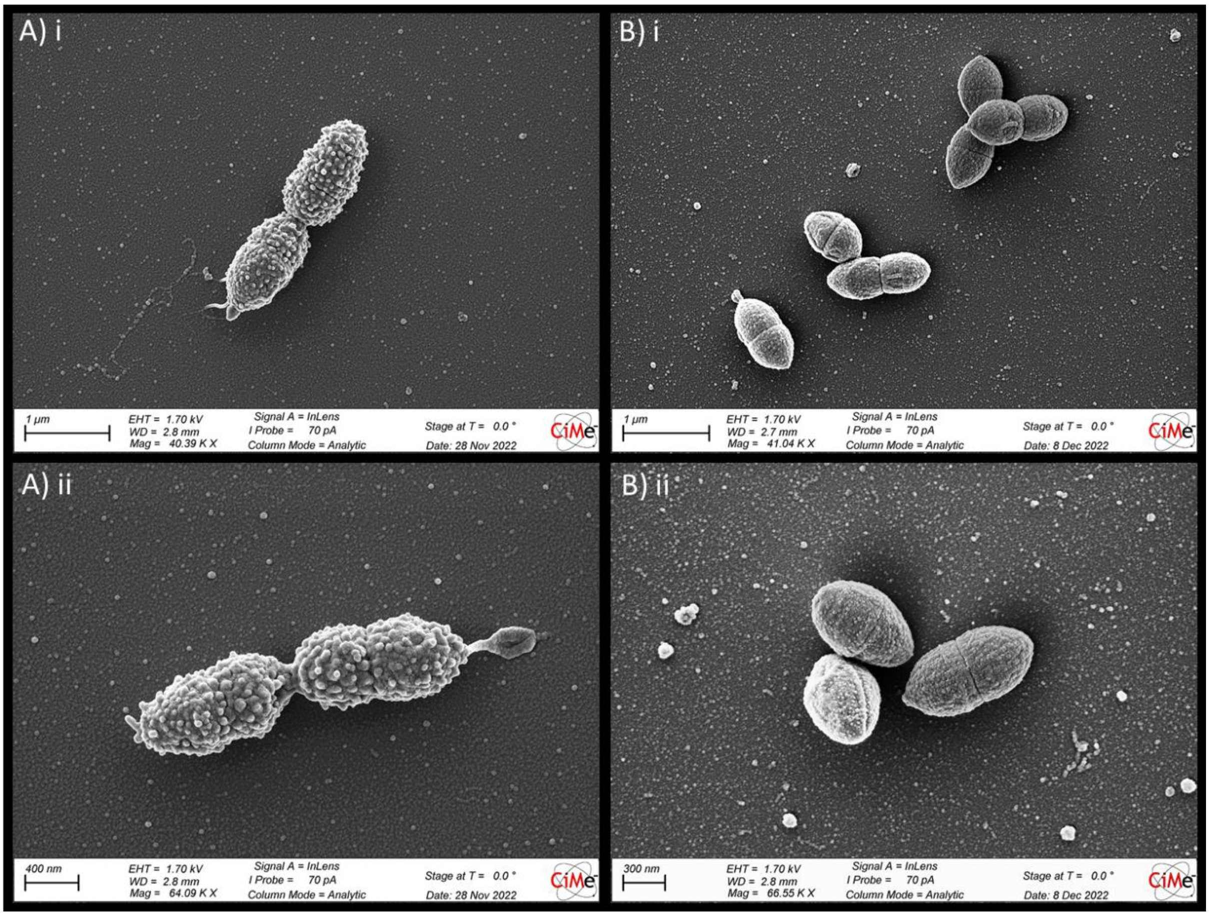


**Supplementary Figure S7** – **A)** Wild-type *Streptococcus pneumoniae* and **B)** a genetically modified non-encapsulated derivative (termed R6) were washed 3× in PBS, then resuspended at $10^{10}$ CFU/mL final concentration in filter-sterilised PBS for fixation and imaging by scanning electron microscopy (SEM). Images are representative of duplicate samples, imaged at approximately 40,000× (i) and 65,000× (ii) magnification.

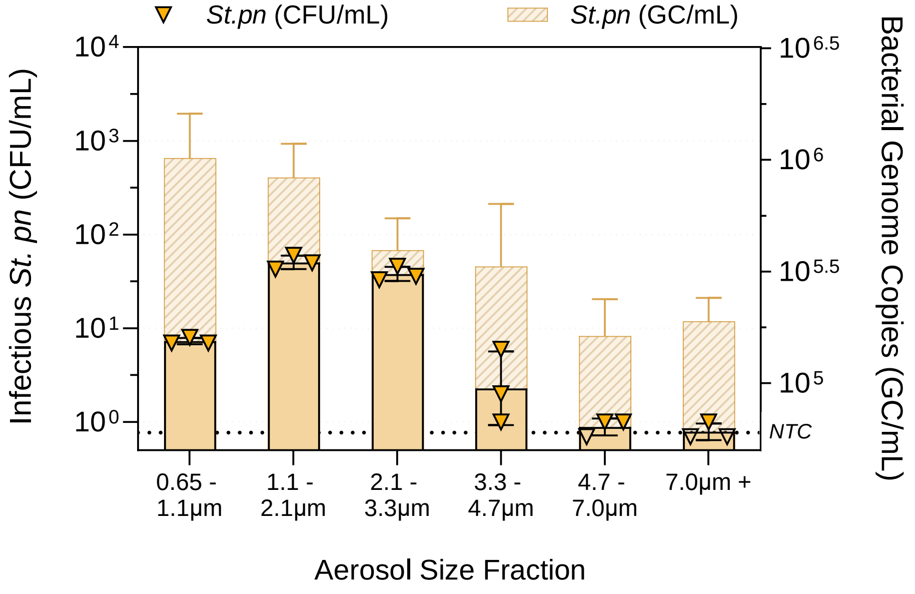


**Supplementary Figure S8** – Recovery of *S. pneumoniae* from aerosol particles at 40% RH. IAV was added to PBS containing live *S. pneumoniae* bacteria at 5 × $10^8$ CFU/mL. Virus was spiked in at 2 × $10^9$ PFU/mL final viral concentration. The liquid inoculum was added to a sparging liquid aerosol generator (SLAG), and nebulised into an aerosol chamber (comprised of a sealed 1.6-$m^3$ polytetrafluoroethylene (PTFE) chamber suspended inside a large biosafety cabinet) for a total of 30 seconds, with the air-flow set at 30L air/min. The chamber was maintained at 24 ± 1°C and at the targeted RH of 40% (± 3%) for the full duration of each experiment. Immediately after the 30 second nebulisation, aerosol particles were recovered using an Andersen Impactor for a total of 15 minutes. Aerosol samples collected by each Stage of the Andersen impactor (6 stages total) were recovered for quantification of infectious *S. pneumoniae* bacteria (CFU/mL, left axis) and bacterial genome copies (GC/mL, right axis). Infectious titers were determined in technical triplicate by agar plating, and bacterial genome copies were determined in technical triplicate by RT-qPCR. Clear symbols indicate samples that were below agar plating limits of quantification (LOQ), and were set a LOQ/√2. NTC indicates the background level obtained for non-template control samples by RT-qPCR. Geometric mean geometric mean ± geometric SD is shown in all cases.

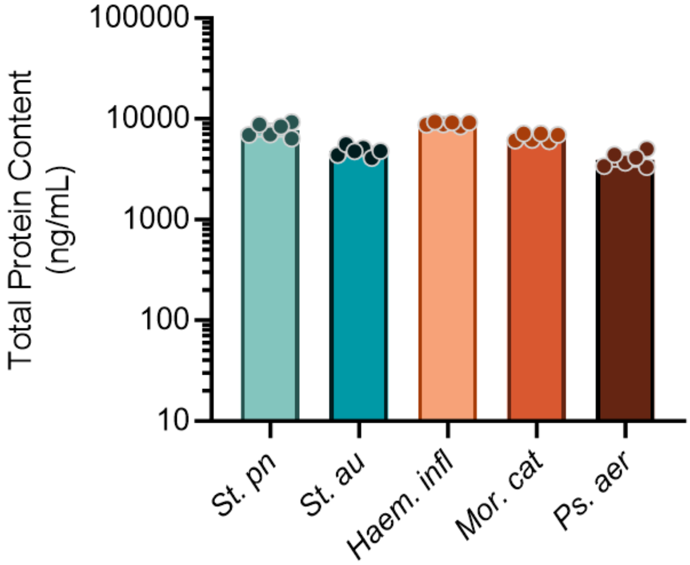


**Supplementary Figure S9 –** Total protein content of respiratory bacterial strains at 1 × $10^8$ CFU/mL or equivalent. All respiratory bacterial strains were grown to comparable early-log phase to generate working stocks, using growth conditions and growth media specified in the Methods section. When required, strains were thawed, washed 3× in PBS to remove residual media, and resuspended at 1 × $10^8$ CFU/mL for use in droplet experiments, with the exception of *M. catarrhalis*. Due to the lower viability of *M. catarrhalis* after freeze-thawing (a known property of this bacterium), diluting based on the viable CFU counts alone resulted in a suspension that was approximately 5× more dense in terms of total protein than all the other prepared strains, due to presence of a higher proportion of dead cells. Thus, to ensure fair comparison to all other strains tested here when mixing with IAV, *M. catarrhalis* was instead diluted to 2 × $10^7$ viable CFU/mL prior to use, equating to 1 × $10^8$ CFU/mL equivalent of total organics. The total protein content in all 5 bacteria suspensions is shown above when stocks were diluted in this comparable manner. For total protein quantification, bacterial pellets were resuspended specifically in miliQ, then heat-inactivated for 20 minutes. Total protein was quantified by Qubit 4 using the Protein Assay Quantification Kit, according to manufacturer's instructions. Data indicates mean protein content (ng/mL) for samples quantified in technical triplicate on 2 separate occasions (n = 6 samples total per bacterial strain).

**For simplicity, all strains are stated as being diluted to 1 × $10^8$ CFU/mL for use in experiments in the Main Text.**
